# A single cell atlas of human and mouse white adipose tissue

**DOI:** 10.1101/2021.11.09.466968

**Authors:** Margo P. Emont, Christopher Jacobs, Adam L. Essene, Deepti Pant, Danielle Tenen, Georgia Colleluori, Angelica Di Vincenzo, Anja M. Jørgensen, Hesam Dashti, Adam Stefek, Elizabeth McGonagle, Sophie Strobel, Samantha Laber, Saaket Agrawal, Gregory P. Westcott, Amrita Kar, Molly L. Veregge, Anton Gulko, Harini Srinivasan, Zachary Kramer, Eleanna De Filippis, Erin Merkel, Jennifer Ducie, Christopher G. Boyd, William Gourash, Anita Courcoulas, Samuel J. Lin, Bernard T. Lee, Donald Morris, Adam Tobias, Amit V. Khera, Melina Claussnitzer, Tune H. Pers, Antonio Giordano, Orr Ashenberg, Aviv Regev, Linus T. Tsai, Evan D. Rosen

**Author notes:** Affiliated with Broad Institute while research was conducted (Samantha Laber). **Address correspondence to:** Evan D. Rosen, MD PhD, Division of Endocrinology, Diabetes, and Metabolism Beth Israel Deaconess Medical Center, 330 Brookline Avenue, Boston, MA 02215.

## Abstract

White adipose tissue (WAT), once regarded as morphologically and functionally bland, is now recognized to be dynamic, plastic, heterogenous, and involved in a wide array of biological processes including energy homeostasis, glucose and lipid handling, blood pressure control, and host defense^1^. High fat feeding and other metabolic stressors cause dramatic changes in adipose morphology, physiology, and cellular composition^1^, and alterations in adiposity are associated with insulin resistance, dyslipidemia, and type 2 diabetes (T2D)^2^. Here, we provide detailed cellular atlases of human and murine subcutaneous and visceral white fat at single cell resolution across a range of body weight. We identify subpopulations of adipocytes, adipose stem and progenitor cells (ASPCs), vascular, and immune cells and demonstrate commonalities and differences across species and dietary conditions. We link specific cell types to increased risk of metabolic disease, and we provide an initial blueprint for a comprehensive set of interactions between individual cell types in the adipose niche in leanness and obesity. These data comprise an extensive resource for the exploration of genes, traits, and cell types in the function of WAT across species, depots, and nutritional conditions.

## A single cell atlas of human white adipose tissue

Mature adipocytes are too large and fragile to withstand traditional single cell approaches; as a result, several groups have focused on the non-adipocyte stromal-vascular fraction (SVF) of mouse^3–6^ and human^7^ adipose tissue. An alternative strategy involves single nucleus (sNuc) sequencing, which can capture adipocytes, and has been used to describe murine epididymal^8, 9^ and human brown adipose tissue^10^. To compare these approaches in the context of human WAT, we pursued experiments on two cohorts of subjects. In the first, we collected subcutaneous WAT from 9 women, isolated single cells from the SVF using collagenase digestion, and then performed whole cell Drop-seq [hereafter referred to as single cell (sc)RNA-seq]. Because different depots have been differentially linked to metabolic disease^11^, for the second cohort we collected paired subcutaneous (SAT) and omental visceral (VAT) adipose tissue from 10 individuals, and SAT alone from three additional individuals (10 women, 3 men), and performed sNuc-seq (**Figures 1a, b**, and **Extended Data Table 1**). Doublet and low-quality filtering left 166,149 total cells (28,465 single cells and 137,684 single nuclei). The data from both approaches were integrated, enabling the identification of the canonical cell types found in WAT, including adipocytes, ASPCs, vascular cells, and immune cells (**Figures 1c, d**; **Supplementary Table 1**). As expected, adipocytes were found only in the sNuc-seq dataset. The sNuc-seq data was also enriched for vascular cells and macrophages, likely because collagenase digestion did not fully dissociate these cell types. Mesothelial cells were not seen in the scRNA-seq dataset, which did not include visceral tissue. Some of the visceral samples included cells that appeared to be endometrial in origin (*PRLR*+), likely due to endometriosis. Overall proportions of adipocytes and ASPCs did not differ between depots, but depot clearly affects the distribution of cells within these populations (**Extended Data Figure 1a, b**, **2a, b**, **Extended Data Table 2**). In our limited cohort, we could not detect major effects of BMI on cell type proportions. To assess this finding at larger scale, we utilized our dataset as a reference to estimate cell type proportions in bulk-RNA sequencing data^12^ obtained from the SAT of 331 men in the METSIM cohort^13^. This deconvolution analysis found that the relative abundance of adipocytes in that cohort was negatively correlated with BMI, while ASPCs and myeloid cells were positively correlated (**Figure 1e**).

**Fig. 1.**
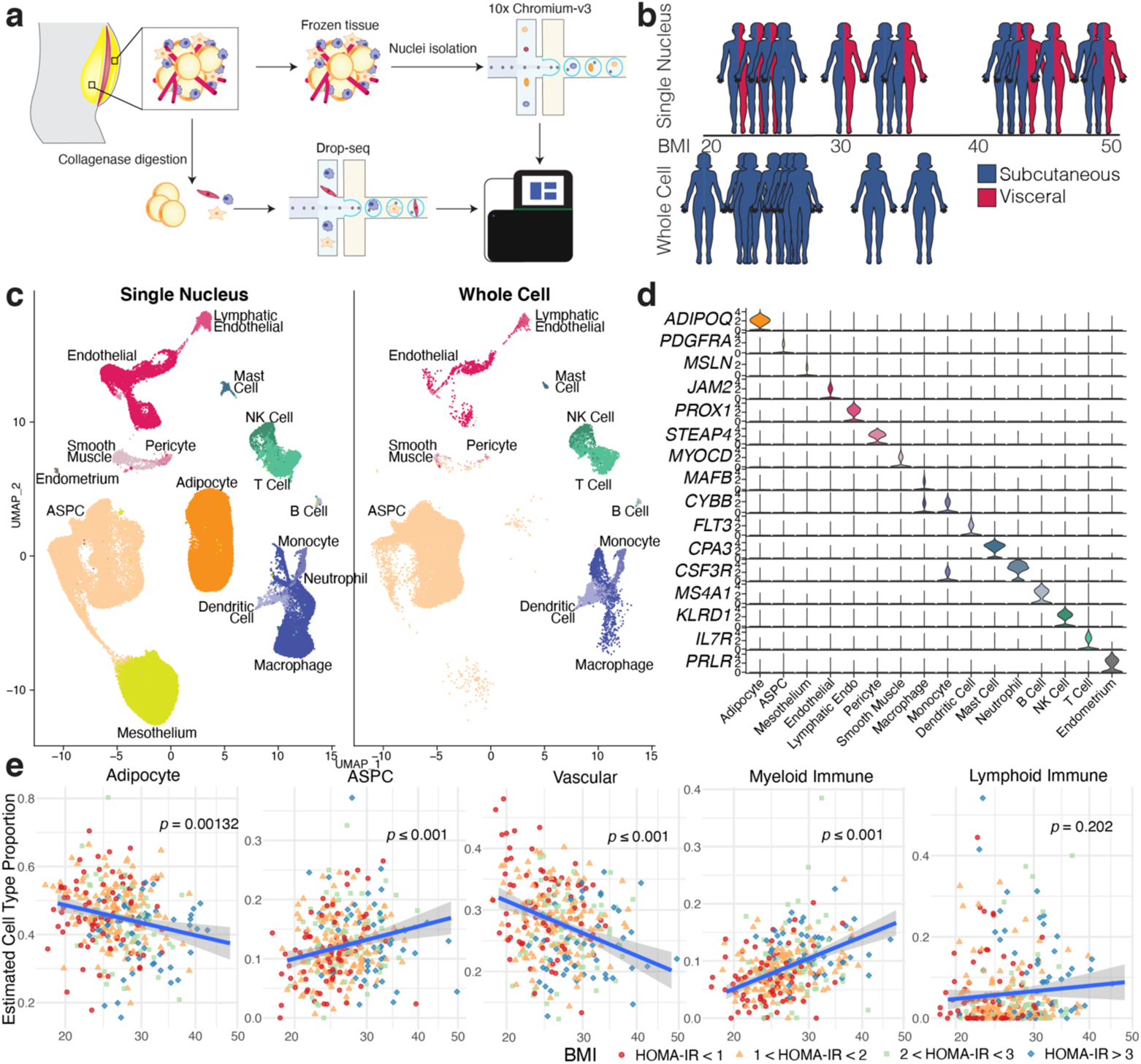
A single cell atlas of human white adipose tissue. **a**, Schematic of workflows for scRNA-seq and sNuc-seq of human WAT. **b,** Graphical representation of the cohorts for both studies. Only the sNuc-seq cohort contains VAT. **c,** UMAP projection of all 166,129 sequenced human cells split by cohort. **d,** Marker genes for each cell population in the human WAT dataset. **e,** Estimated cell type proportions in bulk RNA sequencing data of subcutaneous adipose tissue from 331 individuals from the METSIM cohort calculated using sNuc-seq data as reference. Vascular cells include endothelial, lymphatic endothelial, pericytes, and smooth muscle cells. Myeloid immune includes macrophages, monocytes, dendritic cells, mast cells and neutrophils, and lymphoid immune includes B cells, NK cells, and T cells. For lines of best fit: Adipocytes R^2^ = 0.031, ASPCs R^2^ = 0.034, Vascular R^2^ = 0.076, Myeloid Immune R^2^ = 0.13, Lymphoid Immune R^2^ = 0.0049.

**Fig. 2.**
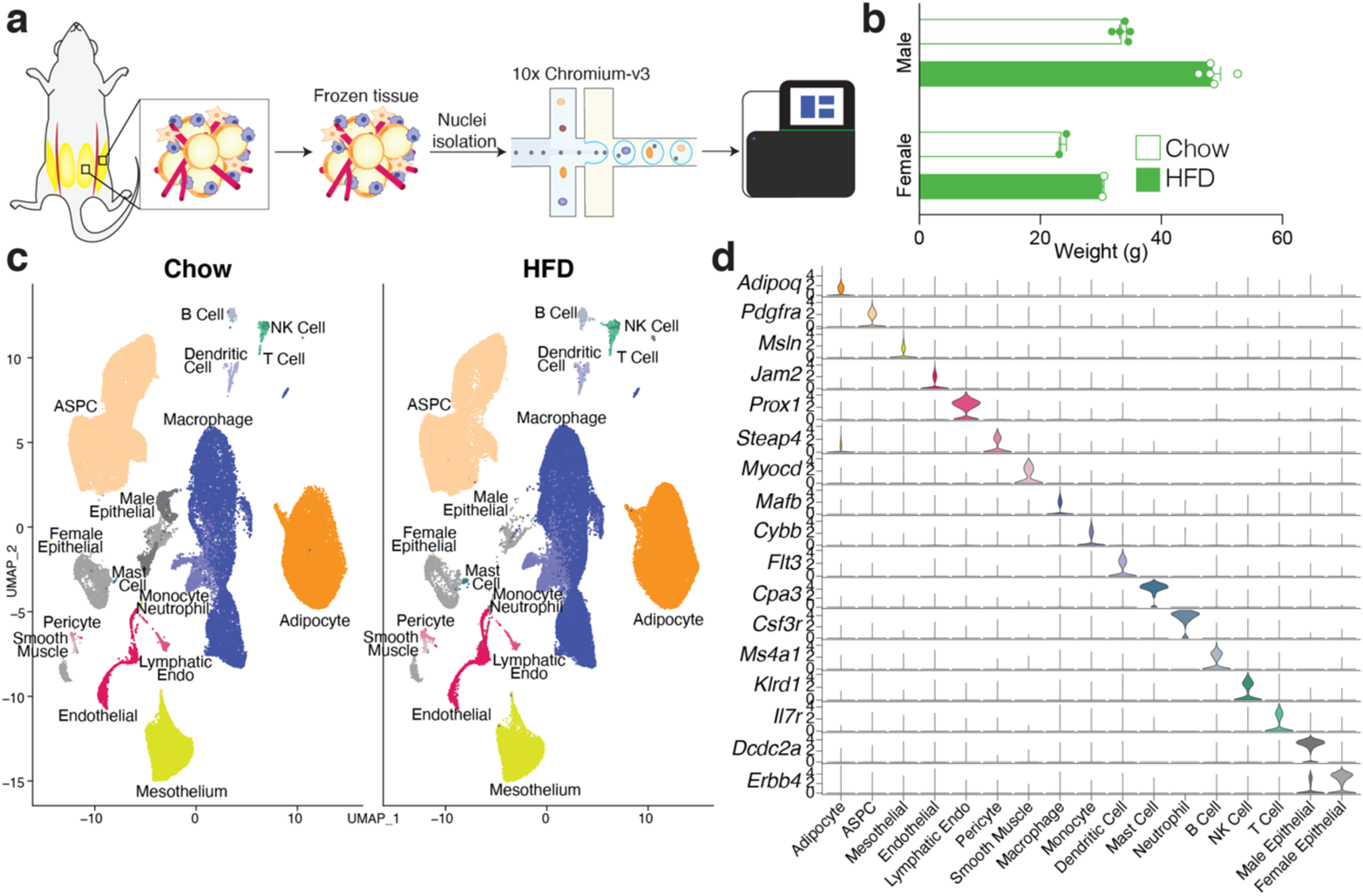
A single cell atlas of mouse white adipose tissue. **a**, Schematic of workflow for sNuc-seq of mouse ING and EPI adipose tissue. **b,** Body weight of chow and high fat fed animals. **c,** UMAP projection of all 197,721 sequenced mouse cells split by diet. **d,** Marker genes for each cell population in the mouse WAT dataset.

## A single cell atlas of mouse white adipose tissue

Murine models are commonly used to study adipose tissue biology^14^. We thus sought to compare mouse and human WAT at the single cell level by performing sNuc-seq on inguinal (ING, corresponding to human SAT) and perigonadal [PG, epididymal (EPI) in males, periovarian (POV) in females, corresponding to human VAT] adipose tissue of mice fed either a chow or high fat diet for 13 weeks (**Figure 2a, b**). After doublet removal and quality filtering, we considered a total of 197,721 cells (106,469 from PG and 91,252 from ING), identifying all cell types observed in human WAT (**Figure 2c, d**; **Supplementary Table 2**) with the addition of distinct male and female epithelial populations (*Dcdc2a*+ and *Erbb4*+, respectively). The female population is largely found in ING samples and resembles mammary epithelial cells, while the male population is almost exclusively found in PG samples, and as noted by others^9^ may represent contaminants from the epididymis and other reproductive structures that are tightly apposed to fat^15^. In contrast to the human data, cell type abundance in mouse WAT are highly dependent on body weight with relatively little variation between depots (**Figure 2c** and **Extended Data Figure 3a, b**, **Extended Data Table 2**). The proportions of cell types in mouse adipose tissue after HFD were notably different between male and female mice, which might reflect a true sex difference, or may reflect that males gain more weight on HFD (**Extended Data Figure 3b**). To compare across species, we used a reference mapping algorithm to assign each mouse cell to a human cluster and noted a high degree of overall similarity between annotated mouse clusters and mapped human clusters (**Extended Data Figure 3c**). Similarly, the proportions of each cell type were roughly similar between humans and chow-fed mice (compare **Extended Data Figure 2b** to **Extended Data Figure 3b**).

**Fig. 3.**
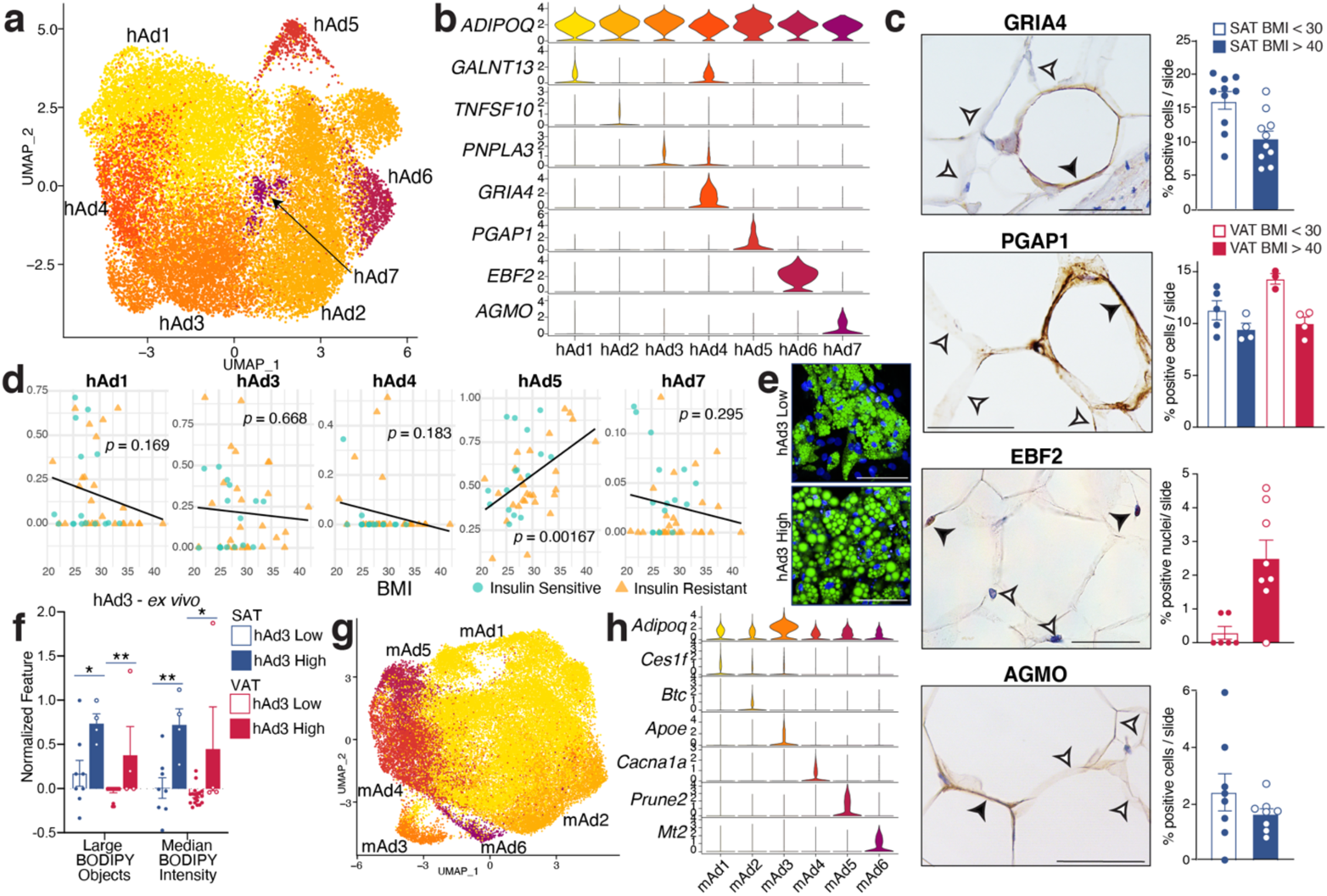
Subclustering of human and mouse adipocytes reveals multiple distinct populations that vary across depot and diet. **a,** UMAP projection of clusters formed by 25,871 human white adipocytes. **b,** Expression of adipocyte marker *ADIPOQ* as well as specific marker genes for each adipocyte subpopulation. **c,** IHC for marker genes of adipocyte subpopulations hAd4, hAd5, hAd6, and hAd7 in human adipose tissue and quantification of percentage of positive adipocytes per slide in lean and obese individuals (GRIA4: 5 lean, 5 obese, 2 slides per person; PGAP1: 5 lean SAT, 4 obese SAT, 3 lean VAT, 4 obese VAT, 1 slide per person; EBF2: 3 lean, 4 obese, 2 slides per person; AGMO: 4 lean, 4 obese, 2 slides per person). Scale bars are 25 μm for GRIA4, EBF2, and AGMO, 20 μm for PGAP1. **d,** Estimated proportions of adipocyte subpopulations in bulk RNA sequencing data of enzymatically isolated subcutaneous adipocytes from 43 individuals plotted against subject BMI. **e,** Representative images of *ex vivo* differentiated human subcutaneous adipocytes predicted to have a low or high amount of hAd3 cells based on deconvolution of bulk RNA sequencing data. Green represents BODIPY staining, blue represents Hoechst staining. Scale bars are 100 μm. **f,** Normalized count of BODIPY-related features in human subcutaneous and visceral adipocytes differentiated *ex vivo* and stratified into low and high hAd3-containing populations. **g,** UMAP projection of clusters formed by 39,934 mouse white adipocytes. **h,** Expression of adipocyte marker *Adipoq* as well as specific marker genes for each mouse adipocyte subpopulation. For bar graphs, error bars represent standard error of the mean (SEM), *, *p* < 0.5, **, *p* < 0.1. For lines of best fit: hAd1 R^2^ = 0.046, hAd3 R^2^ = 0.0045, hAd4 R^2^ = 0.043, hAd5 R^2^ = 0.22, hAd1 R^2^ = 0.027.

## Analysis of human and mouse stromal-vascular cellular subtypes

### Vascular Cells

Subclustering of human vascular cells revealed expected cell types including blood endothelial clusters that represent arteriolar, stalk, and venular cells, as well as lymphatic endothelial cells (LECs), pericytes, and two distinct populations of smooth muscle cells (SMCs) (**Extended Data Figure 4a, b**). Mouse vascular cells formed similar clusters, but with only one SMC cluster (**Extended Data Figure 4c, d**). As expected, reference mapping demonstrated high similarity between human and mouse vascular subclusters (**Extended Data Figure 4e**). The proportions of vascular cells were similar across depots for both mouse and human, although LECs were to be more common in visceral fat of both species (**Extended Data Figure 4f, g**).

**Fig. 4.**
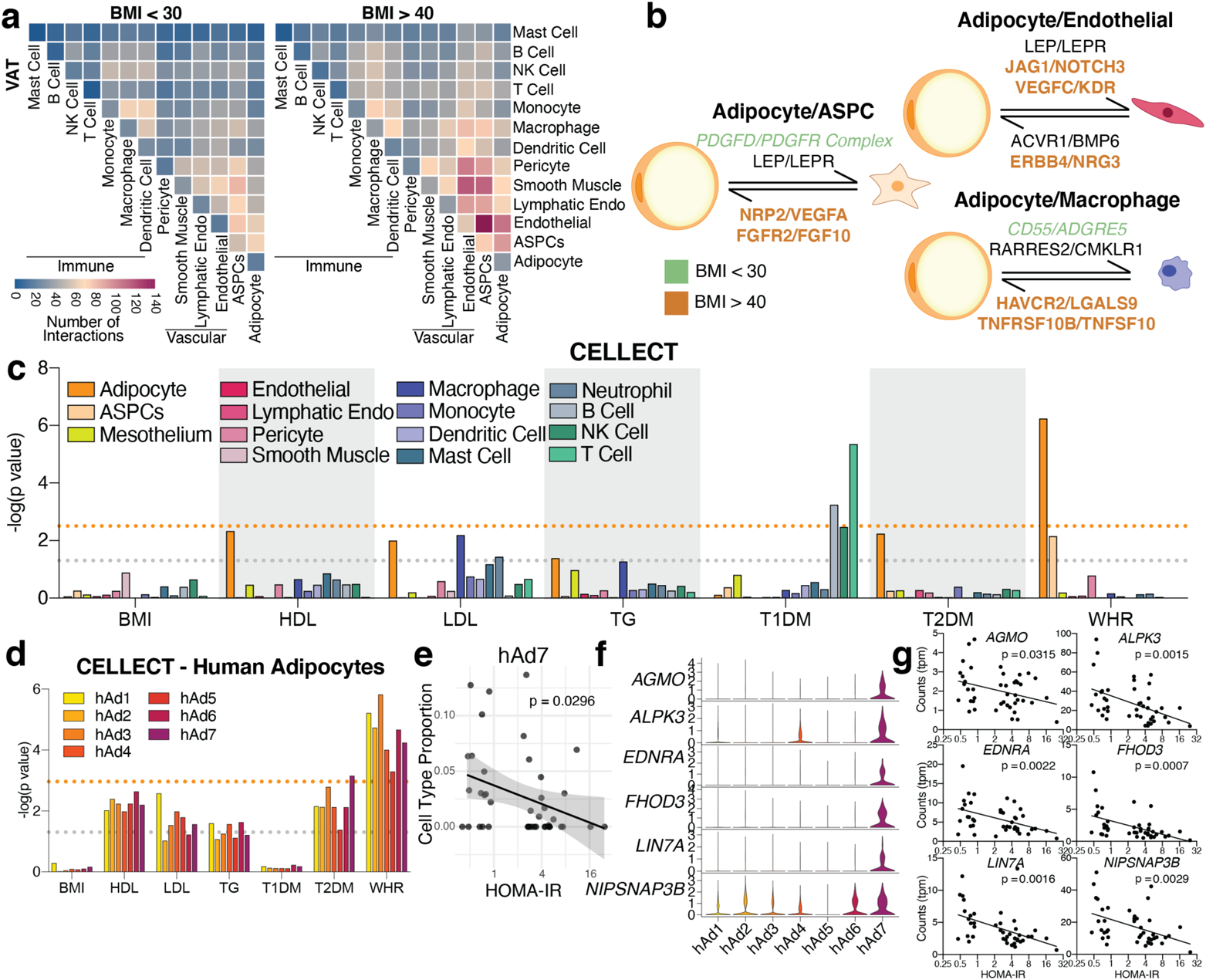
Extensive cell-cell interactions in WAT and associations with human disease traits. **a,** Heatmap showing number of significant interactions identified between cell types in VAT of low (<30) and high (>40) BMI individuals as determined by CellphoneDB. **b,** Selected interactions between adipocytes and ASPCs, endothelial cells, and macrophages identified using CellphoneDB; orange and green indicate interactions that are significant only in BMI > 40 or only in BMI >30, respectively. **c,** CELLECT *p* values of the association between cell types in the human adipose sNuc-seq dataset with GWAS studies. The grey line represents *p* = 0.05 and the orange line represents significant p value after Bonferroni adjustment (*p* = 0.003), based on number of cell types queried. Both T2D and WHR were BMI-adjusted. **d,** CELLECT *p* values for adipocyte subpopulations. The grey line represents *p* = 0.05 and the orange line represents significant *p* value after Bonferroni adjustment (*p* = 0.001), based on all cell subtypes queried. **e,** Estimated cell type proportion of hAd7 in bulk RNA-seq data of enzymatically isolated subcutaneous adipocytes from 43 individuals plotted against HOMA-IR. For line of best fit, R^2^ = 0.11. **f**-**g,** Expression of hAd7 marker genes negatively correlated with HOMA-IR in human adipocyte subpopulations (**f**) and bulk RNA sequencing data of human adipocytes (**g**).

There was little effect of adiposity on vascular cell populations in the human samples; mice, however, showed significant changes in vascular cells after high fat feeding, including a lower proportion of *Dkk2^+^* arteriolar cells and concomitantly higher levels of venular cells. There was also a reduction in the relative proportion of LECs and increased pericytes on HFD (**Extended Data Figure 4f, g**).

### Immune Cells

Analysis of human immune cells from scRNA-seq and sNuc-seq samples again revealed expected cell types, including multiple subpopulations of monocytes, macrophages (*CD14*+), dendritic cells (DCs), B and T lymphocytes, and NK cells (*CD96*+), as well as mast cells (*CPA3*+) and neutrophils (*CSF3R*+) (**Extended Data Figure 5a, b**). These subpopulations resemble known immune cell populations. For example, monocyte subpopulations 1 and 2 resemble classical and non-classical monocytes and DC subpopulations 1 and 2 similarly resemble previously reported *CLEC9A*+ and *CD1C*+ populations from blood, respectively^16^. Lymphocytes also resemble previously reported B cell, T Cell, and NK cell populations from human WAT, including *CTLA4*+ hTregs^17^. Examination of the mouse WAT immune compartment revealed most of the same cell types, although there were notable differences in the relative abundance of myeloid and lymphoid cells between species (**Extended Data Figure 5c, d**). Human WAT contains somewhat fewer T/NK cells than macrophages/monocytes (~30% vs. ~60% of recovered immune cells); this imbalance was greatly exaggerated in murine WAT (macrophages ~90% of recovered immune cells vs. 3% T/NK cells). Because a wealth of data supports a key role for macrophages/monocytes in adipose biology^18, 19^, we separated these cell types from other immune cells *in silico* for subsequent analysis. Mouse clusters of non-monocytes/macrophages mapped relatively well to their human counterparts, with some mixing of T and NK populations (**Extended Data Figure 5e**). Macrophages and monocytes also mapped well to their general class, but this association often broke down when considering macrophage subpopulations (**Extended Data Figure 5f**). Thus, mouse cluster mMac3, which comprises the *Trem2^+^* cells also called “lipid-associated” macrophages^1^ maps well to *TREM2^+^* human hMac2 cells, as expected, but hMac2 also associated with every other mouse macrophage subpopulation, most notably the *Fgf13^+^* mMac1 group (**Extended Data Figure 5f**).

The proportion of immune cell populations was similar in human SAT and VAT, with a few exceptions, such as *PROS1^+^* hMac3 cells which were more abundant in VAT (**Extended Data Figure 6a, e**). In mice, small depot-dependent differences were eclipsed by relatively huge shifts in response to diet in male mice (**Extended Data Figure 6b, d, f**). Most notably, HFD resulted in a massive increase in macrophage numbers, primarily in PG, consistent with a large body of prior data^18, 21^, (**Extended Data Figure 3c, 6f**). As a proportion of total immune cells, HFD induced large shifts in mMac1 (down in ING, up in EPI), mMac2 (down in EPI), and *Trem2^+^* mMac3 (up in ING and EPI) in male mice (**Extended Data Figure 6f**). Reductions in the proportion of most other immune cell types (e.g., NK cells, T and B lymphocytes, DCs, and neutrophils) are likely due to the large influx of macrophages, rather than to intrinsic loss of those specific cell types following HFD (**Extended Data Figure 6b, d, f**). Mast cells increase proportionally after HFD despite the influence of macrophages, as previously reported^22^. Female mice exhibit a much less impressive response to HFD, with the only significantly different diet-related change being a reduction in *Prg4*+ mMac4 cells (**Extended Data Figure 6f**).

Accumulation of adipose tissue macrophages in obesity has also been shown in human WAT, using a combination of histomorphometry and flow sorting^19, 23^. Our data are in general support of this conclusion, though the magnitude of the effect is significantly less prominent than that seen in mouse WAT (**Extended Data 2b, 6c, e, f**). The largest change involves hMac3, which is induced in visceral fat with higher BMI (**Extended Data 6c, e**). We did not observe differential representation of other immune cell in WAT from subjects with high BMI vs. low BMI.

### Mesothelial cells

Subclustering of mesothelial cells revealed three populations in both human VAT and mouse PG (**Extended Data Figure 7a-d**). Only sNuc-seq samples were used in this analysis because our human scRNA-seq data did not include VAT. When mouse mesothelial clusters were mapped to human clusters, cells were split between human clusters hMes1 and hMes2, with no cells mapping to hMes3 (**Extended Data Figure 7e**). The proportions of most mesothelial subpopulations did not vary with obesity or high fat diet, with the exception of hMes1 and hMes2, which were reduced and increased in higher BMIs, respectively. (**Extended Data Figure 7f, g**).

### *ASPCs* (see Supplementary Note 1)

We identified six distinct subpopulations of human ASPCs in subclustered scRNA-seq and sNuc-seq samples, all of which express the common marker gene *PDGFRA* (**Extended Data Figure 8a, b**). Similarly, we noted six subpopulations in the mouse ASPC data, all of which were also *Pdgfra^+^* and some of which correspond well with a particular human subpopulation (**Extended Data Figure 8c-e**). For example, mASPC2 and hASPC2 are both characterized by high expression of *Aldh1a3/ALDH1A3*, and strongly resemble previously identified early multipotent progenitor cells that reside in the reticular interstitium of the fat pad^5^. Similarly, mASPC4 and hASPC4 express *Epha3/EPHA3* and likely represent the anti-adipogenic Areg population reported by Schwalie et. al.^3^. Seeking to better place our mouse ASPC data into the overall context of the published literature, we performed reference mapping between our ASPCs and ASPC populations reported by others^3–6, 9^ and found general agreement across studies (**Extended Data Figure 8f**). As mentioned, mASPC2 cells map to the *Dpp4*+/*Ebf2*+ ASPCs identified by other studies and mASPC1 and mASPC6 map strongly to adipose progenitors, including the *Icam1^+^* cells identified by Merrick et. al.^5^.

Many human and mouse ASPC subclusters showed dependency on diet, depot, or both. hASPC1, hASPC4, and hASPC5 were more prevalent in SAT than VAT, with increases in SAT hASPC4 and hASPC5 proportion in subjects with higher BMI (**Extended Data Figure 9a, c, e**). Conversely, hASPC3 and hASPC6 were more prevalent in VAT. In male mice, early progenitor cells (mASPC2) were notably more abundant in ING than PG; such depot selectivity was not noted for the analogous hASPC2 in humans. mASPC5 and mASPC6 were more prevalent in EPI vs ING, although this varied with obesity (e.g., the proportion of mASPC6 cells was greater in EPI than ING, but only after HFD) (**Extended Data Figure 9b, d, f**). Many of these observations are consistent with previous findings in adipose biology. For example, HFD has been shown to increase adipogenesis specifically in PG in mice^24, 25^. Our data indicates that pre-adipocyte subclusters like mASPC6 increase dramatically in response to HFD in PG only. The loss of early progenitors (mASPC2) in PG with HFD is consistent with conversion of these cells along the differentiative pathway, i.e., toward mASPC6 (**Extended Data Figure 9b, d, f**). These patterns are harder to discern in the human samples, which may reflect the fact that patient data are captured at variable time points after the onset of obesity, whereas the mouse samples are synchronized over a relatively short time period. Nonetheless, we do observe a VAT-specific increase in hASPC6 in subjects with high BMI (BMI > 40) (**Extended Data Figures 8e, 9e**).

## Unique subpopulations of human white adipocytes

White adipocytes are generally considered to be monotypic and essentially uniform in function, although some recent studies have begun to challenge this assumption^8–10, 26^ The high resolution of our data enabled us to find that human white adipocytes cluster into seven subpopulations with distinct markers (**Figure 3a-b**). We noted strong depot-specific associations of adipocyte subtypes, with hAd1, hAd3, hAd4, and hAd7 localized primarily to SAT, while hAd2 and hAd6 were almost exclusively found in VAT. hAd5 represents a smaller population that is roughly equally distributed between SAT and VAT (**Extended Data Figure 10a-c**). We also noted a BMI-dependent shift in adipocyte subtype within both depots (**Extended Data Figure 10b, c**). Importantly, all adipocyte subpopulations are present in the majority of subjects, indicating that these subtype designations are generalizable and do not reflect sample-specific variation (**Extended Data Figure 10c**). Immunohistochemistry (IHC) and/or immunofluorescence of markers for hAd4, hAd5, hAd6, and hAd7 in human subcutaneous or visceral adipose tissue identified specific subpopulations of adipocytes at proportions similar to those seen in the single cell data (**Figure 3c** and **Extended Data Figure 10 d, e**). To examine whether SAT subtype proportion was influenced by BMI in a larger dataset, we estimated individual subtype proportions by deconvolution analysis of bulk RNA-seq data from purified isolated subcutaneous human adipocytes from 43 women (**Figure 3d**). This analysis showed that clusters hAd4 and hAd7 trend to negative correlation with BMI, aligning with our IHC findings, while hAd5 proportion is positively correlated with BMI. Visceral adipocytes are absent from this dataset and so we were unable to assess the prevalence of hAd2 or hAd6 in this cohort, although IHC of hAd6 marker EBF2 also suggests its prevalence may be positively correlated with BMI (**Figure 3c**).

A critical question is whether individual adipocyte subpopulations have specific functions. To assess this, we first looked at genes that participate in the major metabolic activities of adipocytes, including adipokine synthesis and secretion, insulin signaling, lipid handling, and thermogenesis. All subpopulations expressed these genes, although their relative levels differed. Thus, the adipokines adiponectin and adipsin (*CFD*) are most highly expressed in hAd3, and insulin signaling components like *INSR*, *IRS1* and *IRS2* are most highly expressed in hAd5 (**Extended Data Figure 10f**). We next looked more holistically at the data by performing pathway analysis for markers of each subpopulation (**Supplementary Table 3**, **Extended Data Figure 10g-m**). Subpopulations hAd1, which accounts for ~40% of SAT adipocyte nuclei, and hAd2, which accounts for ~60% of VAT adipocyte nuclei, have relatively few specific markers, and the pathways that emerged were similarly unrevealing (**Extended Data Figure 10g, h**). These populations likely represent “basal” subcutaneous or visceral adipocytes, so we therefore focused on subpopulations hAd3-hAd7 for more detailed analysis. hAd3, which comprises ~15% of VAT, was associated with “triglyceride biosynthesis” and included higher expression of *DGAT2*, *SREBF1*, and *PNPLA3* (**Extended Data Figure 10i**). The hAd4 cluster, which makes up ~40% of SAT, expresses the highest levels of several fatty acid desaturases, including *ELOVL5* and *FADS3* (**Extended Data Figure 10j**), which is particularly interesting in light of the insulin-sensitizing role of unsaturated lipokines such as palmitoleate^27^. hAd5 adipocytes comprise a relatively small amount of both SAT and VAT, and besides having the highest expression of several insulin signaling genes, were also characterized by expression of “sphingolipid signaling genes” (**Extended Data Figure 10k**). Both hAd3 and hAd4 express high levels of lipogenic genes, while hAd5 expresses higher levels of lipolysis genes (**Extended Data Figure 10f**).

We next asked whether cultured human adipocytes retain evidence of subpopulation diversity. To that end, we utilized 57 RNA-seq datasets from human subcutaneous and visceral adipocyte progenitors differentiated *ex vivo* over a 14 day timecourse^28^. Deconvolution analysis revealed that many subpopulations identified *in vivo* were retained in the dish. Furthermore, much of the previously noted depot selectivity was recapitulated, such that the visceral subpopulations hAd2 and hAd6 were significantly more likely to appear in cultured visceral cells and the subcutaneous subpopulation hAd4 was overrepresented in cultured subcutaneous cells (**Extended Data Figure 11a**). Furthermore, because these cultured samples were also subjected to high-content image-based profiling using LipocyteProfiler^28^, we were able to correlate individual subpopulations with image-based features representing morphological and cellular phenotypes including lipid and mitochondrial content. Thus, *ex vivo* differentiated adipocyte cultures predicted to have high amounts of hAd3, which express high levels of lipogenic genes and lower levels of lipolytic genes have more overall lipid and larger lipid droplets (**Figure 3e, f**). Conversely, *ex vivo* differentiated adipocyte cultures with high predicted hAd5 content have less overall lipid and smaller lipid droplets, consistent with higher expression of lipolytic genes and less lipogenic gene expression (**Extended Data Figure 11b-d**).

One particularly interesting adipocyte subpopulation is hAd6, which selectively expresses genes typically associated with thermogenesis, such as *EBF2*, *ESRRG*, and *PPARGC1A* (**Extended Data Figure 10l**), a surprising finding given that this population is almost exclusively visceral (**Figure 3c**, **Extended Data Figure 10c**). To better understand the relationship between this subpopulation and visceral adiposity, we looked further into the hAd6 marker EBF2, which has previously been identified as a pro-thermogenic transcription factor^29^. SNPs at the *EBF2* locus are associated with waist-hip ratio (WHR)^30^, which could involve actions in either SAT or VAT. Interestingly, however, a recent study of GWAS loci associated with adiposity in specific depots^31^ found a common variant 15 kb upstream of *EBF2* that was associated specifically with VAT (**Extended Data Figure 12a**). Further analysis revealed that the minor allele of this SNP (MAF = 0.23) was associated with VAT adjusted for BMI and height (VATadj: beta = 0.062 SD per allele, *p* = 1.0 x 10^-12^), but not abdominal subcutaneous (ASAT) or gluteofemoral (GFAT) depots (ASATadj: beta = −0.018 SD per allele, *p* = 0.03), GFATadj: beta = −0.020 SD per allele, *p* = 0.02, **Extended Data Figure 12b**). We additionally stratified individuals into either 0, 1, or 2 carriers of the minor allele and observed an additive trend (G/G median VATadj −0.10 SD, G/A median VATadj = −0.04 SD, A/A median VATadj 0.04 SD; **Extended Data Figure 12c**). Next, we returned to the visceral human adipocytes differentiated *ex vivo*, and found that samples predicted to have a higher proportion of hAd6 adipocytes were characterized by higher mitochondrial intensity and increased expression of mitochondrial and thermogenic genes (**Extended Data Figure 12d-f**). Finally, our analysis of hAd6 markers suggested other pathways associated with thermogenesis, including one for “axon guidance” (**Extended Data Figure 12g**). We could not measure innervation directly using our data, because the nuclei of innervating sympathetic neurons are located in the spinal ganglia and not the fat depot itself. Nonetheless, we estimated relative levels of innervation using the presence of neuron-specific gene expression in the ambient RNA of our visceral sNuc-seq samples. Indeed, the amount of pan-neuronal markers like *TUBB3* (βIII-tubulin) and *UCHL1* (PGP9.5)^32^ strongly correlate with hAd6 proportion (**Extended Data Figure 12e**), further supporting a role for hAd6 as a novel visceral adipocyte subtype with thermogenic potential.

## Adipocytes of mice and humans show critical similarities and differences

Subclustering mouse adipocytes revealed six subpopulations (**Figure 3g, h**). Unlike human adipocytes, mouse adipocyte subtypes exhibit little depot enrichment, especially on chow diet (**Extended Data Figure 13a-c**). There was strong diet-dependency, however, as relative proportions of mAd1 and mAd3 were reduced after HFD, while the opposite was noted for mAd4 and mAd5 (**Extended Data Figure 13b, c**). In contrast to the relatively good cross-species concordance between immune cells, vascular cells, and ASPCs, mouse adipocytes do not map cleanly onto human adipocyte subpopulations. The majority of murine ING adipocytes map most closely to hAd1, while PG adipocytes map to hAd6, with some mapping to hAd2. (**Extended Data Figure 13d-f**).

As in the human, genes associated with major adipocyte functions showed some subpopulation selectivity. For example, lipogenesis genes were highest in HFD-induced population mAd5 (**Extended Data Figure 13c, g**). More detailed pathway analysis on mouse adipocyte subpopulations (**Supplementary Table 3**) showed that the chow-associated clusters mAd1-3 were notably enriched in metabolic pathways, particularly those involved in lipid handling (**Extended Data Figure 13h-j**). The HFD-associated clusters mAd4-6, on the other hand, were linked to pathways like “HIF-1 signaling”, “actin cytoskeleton”, and “NF-κB signaling” (**Extended Data Figure 13k-n**), consistent with the known roles of hypoxia, cytoskeletal remodeling, and inflammation in HFD-induced adipose dysfunction and insulin resistance^23–25^.

Our data allows us to address an important question: are diet-induced changes in gene expression at the population level shared among subpopulations or do they reflect a change in the relative proportion of these subpopulations? To assess this, we examined the twenty most positively and negatively regulated genes from a TRAP-based RNA-seq experiment in white adipocytes from mice fed chow or high fat diet^34^ (**Extended Data Figure 14a**). We noted that some genes, such as *Cyp2e1*, and *Fam13a*, exhibit elevated expression in chow adipocytes in virtually all subpopulations, even those clusters that are selective for HFD (**Extended Data Figure 14b**). However, while the chow-associated gene *Cfd* is reduced in all populations with HFD, expression seems largely driven by the mAd3 population which has the highest expression of *Cfd* and decreases in abundance with HFD (**Extended Data Figure 13b,c**, **14b**). *Sept9*, *Cdkn1a*, and *Fgf13* show increased gene expression after HFD across almost all subpopulations while other HFD-induced genes (e.g., *Slc5a7* and *Dclk1*) increase their expression after HFD in the chow-associated clusters (mAd1-4) but not in the HFD-associated clusters mAd5-7 (**Extended Data Figure 14b**). Thus, diet-dependent expression changes reflect both alterations across all clusters and the emergence or disappearance of distinct populations.

Finally, we were somewhat surprised that we did not see a murine population that could be clearly delineated as thermogenic. Such cells have been noted by others in WAT, even at room temperature^36^. Notably, the distribution of beige adipocytes is not uniform in ING, but tends to be densest close to the inguinal lymph node (LN)^37^. To avoid contamination by LN cells, we excised the node with a fairly wide margin, and it is possible that our samples were thus de-enriched for beige adipocytes. Nonetheless, when we considered the chow fed samples independently, mAd1 split into three clusters (**Extended Data Figure 15a, b**). Two of these clusters, mAd1B and mAd1C, were recognizable as thermogenic beige adipocytes, with relatively high expression of *Prdm16* and *Ppargc1a* in mAd1B and even higher expression of these genes, as well as expression of *Ucp1* and *Cidea* in mAd1C (**Extended Data Figure 15c**). As expected, the thermogenic mAd1B and mAd1C subpopulations were enriched in ING vs. PG samples (**Extended Data Figure 15d, e**) and suggest HFD-induced transcriptional variability masks these subtype designations.

## Exploration of cell-cell interactions within the adipose niche

The functions of WAT are known to be coordinated by neural and hormonal cues from outside the fat pad^38^. There is growing appreciation, however, that intercellular communication within the depot is also critical for the WAT response to overnutrition and other stressors^39^. In particular, attention has focused on cross-talk between adipocytes and immune cells (especially macrophages) in the context of obesity^40^. To assess potential interactions between all identified cell types in different depots and at different body mass, we utilized CellPhoneDB^41^, which utilizes information about the expression of ligand-receptor pairs to estimate cell type communication (**Supplementary Table 4**, **Supplementary Table 5**). As expected, we detected increased potential communication between human adipocytes and macrophages in high BMI vs. low BMI subjects; of 84 potential interactions identified between human adipocytes and macrophages, 40 (48%) were specific for high BMI subjects, while only 3 (4%) were specific for low BMI subjects (**Figure 4a**, **Extended Data Figure 16a, d**). Notably, obesity was also associated with robustly increased expression of genes encoding ligand-receptor pairs between adipocytes and many non-immune cell types, including blood and lymphatic endothelial cells, vascular SMCs, pericytes, and ASPCs (**Figure 4a, b**, **Extended Data Figure 16a, d**). For example, of 145 potential interactions identified between human adipocytes and endothelial cells, 65 (45%) were specific for high BMI subjects, while only 6 (4%) were specific for low BMI subjects (**Extended Data Figure 16d**). Potential interactions between these cell types are frequently bidirectional, and receptors are often expressed on multiple cell types, suggesting networks of communication (**Figure 4b**, **Extended Data Figure 16e**). We also noted differential expression of ligands and receptors within human adipocyte subpopulations, lending further support to the idea that they carry out distinct functions (**Extended Data Figure 16b**). The specific interactions upregulated during obesity suggest that adipocytes play a significant role in obesity-related adipose tissue remodeling. For example, adipocyte expression of angiogenic factors like *JAG1* and *VEGFC* is increased in the obese state, as is true of the expression of their receptors (e.g., *NOTCH3* and *KDR*) on endothelial cells, consistant with obesity-associated induction of angiogenesis by adipocytes^42^ (**Figure 4b**, **Supplementary Table 6**).

Analysis of the mouse data yielded similar results, as HFD increased the intensity of ligand-receptor pair expression, with the most prominent interactions again occurring between non-immune cell types, especially between ASPCs and adipocytes, pericytes, and SMCs (**Extended Data Figure 16c**). Interactions between WAT cell types include several that have been studied, such as the effect of the adipokine leptin on endothelial cells via LEPR^43^, and the actions of TGFB1 on adipose fibrosis via TGFBR1^34^. The majority of these interactions, however, are unstudied in the context of WAT function and dysfunction.

In human samples, most interactions between adipocytes and endothelial cells were shared between SAT and VAT (61%), but of those interactions not shared between depots, the majority were seen in SAT (31% vs. 8% specific for VAT). This same pattern was seen when looking at adipocyte-ASPC interactions (38% SAT-specific vs. 11% VAT-specific), and adipocyte-macrophage interactions (27% SAT-specific vs. 12% VAT-specific). In mice, we noted a more even split between ING- and EPI-specific interactions (e.g., 13% ING-specific vs. 12% EPI-specific adipocyte-endothelial interactions). Adipose niche interactions were only modestly conserved between mouse and human. (**Extended Data Figure 16d**).

## Relationships between WAT cell types and human disease

Adiposity is associated with a wide range of metabolic diseases and traits, and GWAS studies have suggested a specific link between WAT and coronary artery disease (CAD), BMI-adjusted T2D, dyslipidemia, and BMI-adjusted waist-hip ratio (WHR, a measure of body fat distribution)^44–46^. To determine which specific cell types in WAT are likely to mediate these associations, we employed CELLECT, a method for integrating scRNA-seq and sNuc-seq data with GWAS^47^. As expected, Type 1 Diabetes (T1D) was significantly associated with B and T lymphocytes and NK cells, consistent with the known autoimmune basis of that disease (**Figure 4c**). No WAT cell type associated with BMI, as expected given the strong neuronal basis of body weight regulation^48^. The strongest phenotypic association for white adipocytes was with BMI-adjusted WHR, and associations approaching significance were also noted between adipocytes and HDL and T2D (**Figure 4c**, **Supplementary Table 7**).

Because all adipocyte subpopulations were significantly associated with WHR (**Figure 4d**), we looked for adipocyte genes responsible for the association with WHR that are not specific to any particular subpopulation. One such gene is *PPARG*, which is highly expressed in all adipocytes (**Extended Data Figure 17a**). Data from the METSIM cohort indicates a strong inverse relationship between WHR and *PPARG* levels in whole WAT (**Extended Data Figure 17b**). Unfortunately, WHR was not recorded in the cohort used to generate our floated human adipocytes. WHR is, however, highly correlated with HOMA-IR^11^, and we found that PPARG levels showed a strong inverse relationship with HOMA-IR in both the METSIM cohort and in our floated adipocytes (**Extended Data Figure 17c, d**). Furthermore, SNPs in the *PPARG* gene that are associated with BMI-adjusted WHR^30^ are also significantly associated with *PPARG* mRNA levels and HOMA-IR in our floated adipocyte cohort (**Extended Data Figure 17e-h**).

Adipocytes were also the cell type most likely to mediate the association of WAT with T2D, with the strongest association specifically with hAd7 (**Figure 4d**). To further investigate the association between hAd7 and T2D, we took our deconvolved bulk RNA-seq data from floated human adipocytes and plotted the abundance of hAd7 as a function of HOMA-IR. This revealed that hAd7 shows significant inverse correlation with insulin resistance (**Figure 4e**). We then searched for specific hAd7 marker genes that exhibit this same relationship with HOMA-IR, and identified several, including *AGMO*, *ALPK3*, *FHOD3,* and *LIN7A* (**Figure 4f, g**). Of note, *AGMO* (also called *TMEM195*) has emerged as a candidate locus in T2D GWAS^49, 50^.Taken together, our data suggest that hAd7 may have an outsized influence on the risk of T2D, despite representing only ~1% of human adipocytes.

Additionally, although adipocytes did not meet genome-wide significance for an association with LDL, we were struck by the near significant relationship between LDL and hAd1, and to a lesser extent, hAd4 (**Figure 4c, d**). We noted several genes that were selective for hAd1 and/ hAd4, including *NRCAM*, *PEMT*, *PCDH7*, and *VGLL3*, all of which showed a strong positive relationship between expression and LDL levels in our floated adipocyte cohort (**Extended Data Figure 17i, j**)

We also performed CELLECT using the mouse data and noted associations between BMI-adjusted WHR and murine adipocytes (particularly mAd1, mAd3, and mAd6), as well as pre-adipocytes (especially mASPC2) (**Extended Data Figure 18a-c**). This suggests that WHR may be determined in large part by alterations in adipocyte differentiation, a hypothesis consistent with the *PPARG* data above, and with independent studies of different WHR genes^51^. HDL and TG levels are also associated with mouse white adipocyte gene expression (**Extended Data Figure 18a-c**).

## Discussion

Here, we present a comprehensive atlas of human and mouse WAT across depot and nutritional state. Our analysis reveals a rich array of cell types, including blood and lymphatic vascular cells, immune cells, and ASPCs, in addition to adipocytes. These cell types are grossly similar across species, but differ more profoundly when cellular subpopulations are explored. It is tempting to attribute these subpopulation differences to divergence across 65 million years of evolution, but other factors also need to be considered. For example, the human samples were collected after a fast, while the mice were harvested after *ad libitum* feeding, which might be expected to cause some differences in cell state related to insulin signaling or related pathways. Ongoing studies are focused on addressing potential effects of fasting/feeding on WAT composition.

Our dataset reveals subpopulations of human white adipocytes that are associated with a range of adipocyte functions, from lipolysis and lipogenesis to thermogenesis, as well as with phenotypes such as BMI, WHR, and T2D. The single cell resolution of our dataset enables the identification of heterogeneity that cannot be appreciated by bulk RNA sequencing, such as a potentially visceral thermogenic subpopulation (hAd6), and a rare subpopulation associated with T2DM (hAd7). Our dataset provides a rich resource to identify other disease-associated cell types and to better interpret GWAS studies of metabolic phenotypes.

Overall, our data highlight a central role for adipocytes in the local regulation of the adipose depot as well as in systemic physiology. We additionally provide a framework for mouse-human comparison in studies of adipose tissue that will be an important resource for groups hoping to translate murine findings to human treatments. These data provide a lens of unprecedented acuity that better informs our understanding of WAT biology and enables a deeper exploration of the role of adipose tissue in health and disease.

## Supporting information

Supplemental guide and note

Supplementary Table 1 Human Markers

Supplementary Table 2 Mouse Markers

Supplementary Table 3 Adipocyte Cluster Pathway Analysis

Supplementary Table 4 CellphoneDB p values

Supplementary Table 5 Avarage Expression for CellphoneDB

Supplementary Table 6 CellphoneDB Significant Interactions

Supplementary Table 7 CELLECT Results

## METHODS

### Collection of human adipose tissue samples

#### Drop-Seq and Floated adipocyte bulk RNA-seq

Subcutaneous adipose tissue was collected under Beth Israel Deaconess Medical Center Committee on Clinical Investigations IRB 2011P000079. Potential subjects were recruited in a consecutive fashion, as scheduling permitted, from the plastic surgery operating room rosters at Beth Israel Deaconess Medical Center. Male and female subjects over the age of 18 undergoing elective plastic surgery procedures and free of other acute medical conditions were included and provided written informed consent preoperatively. Excess adipose tissue from the surgical site was collected at the discretion of the surgeon during the normal course of the procedure. Subjects with a diagnosis of diabetes, or taking insulin-sensitizing medications such as thiazolidinediones or metformin, chromatin-modifying enzymes such as valproic acid, anti-retroviral medications, or drugs known to induce insulin resistance such as mTOR inhibitors or systemic steroid medications, were excluded.

#### sNuc-Seq

Subcutaneous and visceral adipose tissue was collected under BIDMC Committee on Clinical Investigations IRB 2011P000079 and University of Pittsburgh Medical Center STUDY 19010309. At BIDMC, potential subjects were recruited in a consecutive fashion, as scheduling permitted, from the gynecological, vascular, and general surgery rosters. Male and female subjects over the age of 18 undergoing plastic surgery (panniculectomy, thighplasty or deep inferior epigastric perforators), gynecological surgery (total abdominal hysterectomy and bilateral salpingo-oophorectomy) or general surgery (cholecystectomy (CCY) or colin polyp surgery) and free of other acute medical conditions were included and provided written informed consent preoperatively. Excess adipose tissue from the surgical site was collected at the discretion of the surgeon during the normal course of the procedure. The exclusion criteria were any subjects taking thiazolidinediones, chromatin-modifying enzymes such as valproic acid, anti-retroviral medications, and drugs known to induce insulin resistance such as mTOR inhibitors or systemic steroid medications. At UPMC, inclusion criteria were patients receiving bariatric surgery (Vertical Sleeve Gastrectomy or Roux en Y Gastric Bypass) or lean controls (hernia or CCY surgeries) ages 21-60, exclusion criteria were diagnosis of diabetes (Type 1 or Type 2), pregnancy, alcohol or drug addiction, bleeding or clotting abnormality, or inflammatory abdominal disease. All patients provided written informed consent preoperatively. Excess adipose tissue from the surgical site was collected at the discretion of the surgeon during the normal course of the procedure. 200-500 mg samples were flash frozen immediately after collection for downstream processing.

### Mouse adipose tissue samples

All animal experiments were performed under a protocol approved by the BIDMC Institutional Animal Care and Use Committee. Male C57Bl/6J 16-week-old high fat diet fed (JAX 380050) and chow fed (JAX 380056) mice were obtained from The Jackson Laboratory and maintained on 60% high fat diet (Research Diets, D12492) or chow diet (8664 Harlan Teklad, 6.4% wt/wt fat), respectively, for three weeks before sacrifice. Female 6-week-old chow fed C57Bl/6J mice (JAX 380056) were maintained on 60% high fat diet for 13 weeks before sacrifice. Mice were maintained under a 12 hr light/12hr dark cycle at constant temperature (23°C) with free access to food and water.

### Mature human adipocyte sample preparation

#### Purification of mature human adipocytes

Whole tissue subcutaneous adipose specimens were freshly collected from the operating room. Skin was removed, and adipose tissue was cut into 1- to 2-inch pieces and rinsed thoroughly with 37°C PBS to remove blood. Cleaned adipose tissue pieces were quickly minced in an electric grinder with 3/16-inch hole plate, and 400 ml of sample was placed in a 2-l wide-mouthed Erlenmeyer culture flask with 100 ml of freshly prepared blendzyme (Roche Liberase TM, research grade, cat. no. 05401127001, in PBS, at a ratio of 6.25 mg per 50 ml) and shaken in a 37 °C shaking incubator at 120 r.p.m. for 15–20 min to digest until the sample appeared uniform. Digestion was stopped with 100 ml of freshly made KRB (5.5 mM glucose, 137 mM NaCl, 15 mM HEPES, 5 mM KCl, 1.25 mM CaCl2, 0.44 mM KH2PO4, 0.34 mM Na2HPO4 and 0.8 mM MgSO4), supplemented with 2% BSA. Digested tissue was filtered through a 300 μM sieve and washed with KRB/albumin and flow through until only connective tissue remained. Samples were centrifuged at 233g for 5 min at room temperature, clear lipid was later removed, and floated adipocyte supernatant was collected, divided into aliquots and flash-frozen in liquid nitrogen.

#### Sample selection and Bulk-RNA-seq library construction

Fasting serum was collected and insulin, glucose, free fatty acids, and a lipid panel were measured by Labcorp. BMI measures were derived from electronic medical records and confirmed by self-reporting, and measures of insulin resistance, the homeostasis model assessment-estimated insulin resistance index (HOMA-IR) and revised quantitative insulin sensitivity check index (QUICKI) were calculated^52, 53^. Female subjects in the first and fourth quartiles for either HOMA-IR or QUICKI and matched for age and BMI were processed for RNA-seq.

Total RNA from ~400 μl of thawed floated adipocytes was isolated in TRIzol reagent (Invitrogen) according to the manufacturer’s instructions. For RNA-seq library construction, mRNA was purified from 100 ng of total RNA by using a Ribo-Zero rRNA removal kit (Epicentre) to deplete ribosomal RNA and convert into double-stranded complementary DNA by using an NEBNext mRNA Second Strand Synthesis Module (E6111L). cDNA was subsequently tagmented and amplified for 12 cycles by using a Nextera XT DNA Library Preparation Kit (Illumina FC-131). Sequencing libraries were analyzed with Qubit and Agilent Bioanalyzer, pooled at a final loading concentration of 1.8 pM and sequenced on a NextSeq500.

### Single Cell and Single Nucleus sample preparation and processing

#### SVF isolation and Drop-seq

Adipose tissue samples were collected and processed as above. After removal of floated adipocytes, remaining supernatant was aspirated and the remaining pelleted stromal vascular fraction (SVF)was combined from multiple tubes. The combined SVF was washed 2 times with 50ml cold PBS with 233g for 5 min centrifugation between washes. Erythrocytes were depleted with two rounds of 25 ml. ACK lysing buffer (Gibco™ A1049201) exposure (5 minutes at RT followed by 233g x 5 min centrifugation). Remaining SVF pellet was further washed x 2 with 50ml cold PBS prior to counting on hematocytometer and loading onto Drop-seq microfluidic devices. Drop-seq was performed as described^54^, with the following modifications: first, flow rates of 2.1 mL/h were used for each aqueous suspension and 12 mL/h for the oil. Second, libraries were sequenced on the Illumina NextSeq500, using between 1.6-1.7 pM in a volume of 1.2 mL HT1 and 3 mL of 0.3 μM Read1CustSeqB (GCCTGTCCGCGGAAGCAGTGGTATCAACGCAGAGTAC) using 20 x 8 x 60 read structure.

#### sNuc-Seq

Nuclei were isolated from frozen mouse and human adipose tissue samples for 10x snRNA-seq using a slightly modified approach to what was previously described^55–57^. Samples were kept frozen on dry ice until immediately before nuclei isolation, and all sample handling steps were performed on ice. Each flash-frozen adipose tissue sample was placed into a gentleMACS C tube (Miltenyi Biotec) with 2 mL freshly prepared TST buffer (0.03% Tween 20 [Bio-Rad], 0.01% Molecular Grade BSA [New England Biolabs], 146 mM NaCl [ThermoFisher Scientific], 1 mM CaCl2 [VWR International], 21 mM MgCl2 [Sigma Aldrich], and 10 mM Tris-Hcl pH 7.5 [ThermoFisher Scientific] in Ultrapure water [ThermoFisher Scientific]) with or without 0.2 U/µL of Protector RNase Inhibitor (Sigma Aldrich). gentleMACS C tubes were then placed on the gentleMACS Dissociator (Miltenyi Biotec) and tissue was dissociated by running the program “mr_adipose_01” twice, and then incubated on ice for 10 minutes. Lysate was passed through a 40 µm nylon filter (CellTreat) and collected into a 50 mL conical tube (Corning). Filter was rinsed with 3 mL of freshly prepared ST buffer buffer (146 mM NaCl, 1 mM CaCl2, 21 mM MgCl2; 10 mM Tris-Hcl pH 7.5) with or without 0.2 U/µL RNase Inhibitor, and collected into the same tube. Flow-through was centrifuged at 500 x g for 5 minutes at 4°C with brake set to low. Following centrifugation, supernatant was removed, and the nuclear pellet was resuspended in 50 - 200 µl PBS pH 7.4 (ThermoFisher Scientific) with 0.02% BSA, with or without 0.2U/µL RNase Inhibitor. In order to reduce ambient mRNA, the nuclear pellets of some samples were washed 1-3 times with 5 mL of PBS-0.02% BSA before final resuspension. An aliquot of nuclei from each sample was stained with NucBlue (Thermofisher Scientific), counted in a hemocytometer using fluorescence to identify intact nuclei, and then immediately loaded on the 10x Chromium controller (10x Genomics) according to the manufacturer’s protocol.

For each sample, 10,000-16,500 nuclei were loaded in one channel of a Chromium Chip (10x Genomics). The Single Cell 3’ v3.1 chemistry was used to process all samples. cDNA and gene expression libraries were generated according to the manufacturer’s instructions (10x Genomics). cDNA and gene expression library fragment sizes were assessed with a DNA High Sensitivity Bioanalyzer Chip (Agilent). cDNA and gene expression libraries were quantified using the Qubit dsDNA High Sensitivity assay kit (ThermoFisher Scientific). Gene expression libraries were multiplexed and sequenced on the Nextseq 500 (Illumina) with a 75-cycle kit and the following read structure: Read 1: 28 cycles, Read 2: 55 cycles, Index Read 1: 8 cycles.

### Sequencing, read alignments, and quality control

#### Single-cell/nucleus RNA-seq data analysis

Raw sequencing reads were demultiplexed to FASTQ format files using bcl2fastq (Illumina; version 2.20.0). Digital expression matrices were generated from the FASTQ files using the Drop-Seq tools (https://github.com/broadinstitute/Drop-seq) pipeline, with appropriate adjustments made to the default program parameters to account for the different read-structures in the scRNA Drop-Seq data and sNuc 10X data. Reads from mouse and human were aligned with STAR^58^ (version 2.7.3) against the GRCm38 and GRCh38 genome assemblies, respectively. Gene counts were obtained, per-droplet, by summarizing the unique read alignments across exons and introns in appropriate GENCODE annotations (release 16 of the mouse annotation and release 27 of the human annotation). In order to adjust for downstream effects of ambient RNA expression within mouse nuclei (hereafter “cells”), we used CellBender^59^ (version 0.2.0) to remove counts due to ambient RNA molecules from the count matrices and to estimate the true cells. We also used CellBender to distinguish droplets containing cells from droplets containing only ambient RNA, by selecting droplets with >50% posterior probability of containing a cell. We compared the true cell estimation obtained using CellBender against the same using the DropletUtils software package^60^, which estimates ambient RNA expression levels but does not remove any ambient counts, keeping only the cells that were marked as not ambient by both algorithms. To address ambient RNA in the human sNuc data, we calculated spliced and unspliced RNA content in each cell, because nuclei have a high unspliced RNA content, a high percentage of spliced RNA indicates a high ambient RNA content. We therefore removed sNuc-seq cells containing over 75% spliced RNA. All samples were assessed for doublet content using scrublet^61^ version 0.2.1, and cells called as doublets were removed before further analysis. All cells were further filtered to have greater than 400 UMIs with <10% of UMIs from mitochondrial genes. Genes were filtered such that only genes detected in two or more cells were retained. For the human data, the median number of UMIs detected per cell was 2559 and the median number of genes detected per cell was 1524. For the mouse data, the median number of UMIs detected per cell was 2291 and the median number of genes detected per cell was 1369.

#### Bulk RNA-seq Analysis

Raw sequencing reads were demultiplexed by using bcl2fastq (Illumina). Salmon^62^ (version 1.1.0) was used to simultaneously map and quantify transcript abundances of hg19 genes annotated by release 19 of the GENCODE project’s human reference. Salmon was run using “full” selective alignment (SAF) with mapping validation as described previously^63^. Gene counts were summarized from transcript abundances using the “tximport” package for R^64^.

### Integration, clustering, subclustering, and annotation

Integration, clustering and subclustering analysis were performed using Seurat 3.9.9^65^. The gene counts were normalized using SCTransform^66^, and regressed on mitochondrial read percentage, ribosomal read percentage, and cell cycle score as determined by Seurat. In order to avoid smoothing over depot differences, for integration human and mouse data were grouped by ‘individual’, i.e., if both subcutaneous and visceral adipose tissue for an individual human or mouse were available, they were pooled together during this step. Individuals were integrated with reciprocal PCA, using individuals that had both subcutaneous and visceral samples as references. As a result, the human and mouse references were comprised exclusively from the sNuc seq cohort. To integrate, references were integrated together, then the remaining samples— sNuc seq individuals with only subcutaneous data as well as all Drop-seq samples—were mapped to the reference. For clustering, 5000 variable genes were used, and ribosomal and mitochondrial genes were removed from the variable gene set before running PCA and calculating clusters using a Louvain algorithm, 40 PCs, and a resolution of 0.5. Clusters were identified as adipocytes, preadipocytes, mesothelial cells, vascular cells, or immune cells using marker genes, subset into individual objects, and re-integrated using the above method. Samples with fewer than 50 cells in the subset were removed before re-integration. This led to samples having artificially fewer cells in some instances—for example some Drop-seq samples had cells that clustered with adipocytes, but these cells were removed in subclustering because the small numbers of cells introduced too much variability into the integration. Subclustering was performed using a range of variable genes (1000-2000), PCs (10-40) and resolutions (0.2-0.6).

Markers were calculated using a non-parametric Wilcoxon rank sum test and clusters were evaluated based on the distinctness of called markers to determine the final subclustering conditions. In the subclustered objects, we removed clusters that appeared to represent doublets based on the score assigned by scrublet^61^, or that appeared to be driven by high ambient RNA content as determined by levels of mitochondrial genes and spliced/unspliced RNA ratio. The remaining clusters were annotated based on marker gene expression. In some cases, smaller subclusters (T and NK cells, B cells, monocytes/neutrophils) were further subset and PCA and clustering analysis but not integration was re-run in order to assign clusters. After subcluster annotation, identities were mapped back onto the original object and cells that were removed from the subclustered objects were similarly removed from the all-cell object.

### Deconvolution of bulk RNA-seq data

Bulk RNA sequencing data for subcutaneous adipose tissue from the METSIM cohort were obtained as described previously^13^. Only individuals with available metabolic phenotyping data were used for the deconvolution analysis. Bulk RNA sequencing data for floated human adipocytes were obtained described above. Deconvolution analysis was performed using MuSiC^12^ (version 0.1.1) with human sNuc subcutaneous all cell or adipocyte data as reference. Marker genes used for deconvolution can be found in **Supplemental Table 1**.

### Comparison between mouse and human datasets

Mapping of mouse cells onto human clusters was performed using Seurat multimodal reference mapping^67^. To run, for the all-cell and each subset, the mouse data was prepared by extracting the counts matrix from the mouse sNuc object and mapping the mouse gene names to their human orthologs using a database of ortholog mappings from Mouse Genome Informatics (http://www.informatics.jax.org/homology.shtml). In the case of multi-mapping, the first ortholog pair was used. The mouse object was then split by sample and mapped onto the sNuc-seq data from the matching human all-cell or subset object using the RNA assay and PCA reduction.

### Immunohistochemistry

Subcutaneous (abdominal) and omental adipose tissue biopsies belonging to lean and obese women (GRIA4: subcutaneous, 5 lean and 5 obese individuals; PGAP1: subcutaneous, 5 lean, 4 obese, visceral 3 lean, 4 obese; EBF2: omental, 3 lean and 4 obese individuals; AGMO: subcutaneous, 4 lean and 4 obese individuals, for all experiments two slides per individual for GRIA4, EBF2, AGMO, one slide per individual for PGAP1) were fixed (overnight in 4% paraformaldehyde at 4°C, dehydrated, paraffin embedded and sectioned (4μm thick). The following primary antibodies and respective dilution were used: GRIA4, 1:200, Cat #23350-1-AP, Proteintech; PGAP1, 1:400, Cat. #55392-1-AP, Proteintech EBF2, 1:1000, Cat. #AF7006, R&D systems; AGMO (TMEM195) 1:100, Cat #orb395684, Biorbyt. In brief, after rinsing in PBS, tissue slices were blocked with 3% normal goat serum and incubated with the primary antibody in PBS, overnight at 4°C. After a thorough rinse in PBS, sections were incubated in 1:200 v/v biotinylated secondary antibody solution for 30 minutes (Invitrogen), rinsed in PBS and incubated in avidin-biotin-peroxidase complex (ABC Standard, Vector Laboratories), washed several times in PBS and lastly incubated in 3,3′-diaminobenzidine tetrahydrochloride (0.05% in 0.05 M Tris with 0.03% H_2_O_2_; 5 min). After immunohistochemical staining, sections were counterstained with hematoxylin, dehydrated in ethanol, cleared in xylene and covered with coverslip using Eukitt (Merck). All observations were performed using Nikon Eclipse E800 light microscope.

### Immunofluorescence microscopy of mature human adipocytes

Adipocyte immunofluorescence protocol was adapted from Sárvári et al^9^. Abdominal subcutaneous adipose tissue was collected from two adult female human subjects (BMI 24.9 and 40.3) as above and placed on ice. Tissue was minced and digested with 1 mg/mL type II collagenase (Sigma-Aldrich, C6885) in Hanks’ balanced salt solution supplemented with 0.5% fatty acid-free BSA (Sigma-Aldrich, A6003) at 37° in a water bath with constant shaking at 250 rpm. The cell suspension was filtered through a 250 µM nylon mesh strainer (Thermo, 87791) and washed three times with Krebs-Ringer bicarbonate buffer containing 1% fatty acid-free BSA. All washes throughout this protocol were performed without centrifugation to minimize adipocyte damage and loss; cell suspension was maintained upright for at least 5 minutes to allow mature adipocytes to float, and infranatant was removed with a needle and syringe. The floating adipocytes were fixed with 2% PFA and 1% sucrose in PBS for 30 minutes with constant rotation followed by three washes with 2% fatty acid-free BSA in PBS. Adipocytes were subsequently permeabilized with 0.5% Triton-X (Thermo, 28314) in PBS for five minutes, and incubated with 2.5 µg/mL trypsin (Corning, 25053CI) in PBS for 10 minutes at 37° in a water bath with constant shaking. Adipocytes were then blocked with 2% fatty acid-free BSA in PBS for 30 minutes, and incubated overnight at room temperature with rabbit polyclonal anti-GRIA4 (Proteintech, 23350-1-AP) diluted 1:100 in 500 µL 2% fatty acid-free BSA in PBS with constant rotation. The adipocytes were then washed twice for 10 minutes each with 0.1% fatty acid-free BSA and 0.05% Tween-20 (Sigma-Aldrich, P9416) in PBS, followed by incubation with goat anti-rabbit Alexa Fluor 546 (Thermo, A-11035) secondary antibody diluted 1:500 in 2% fatty acid-free BSA for 2 hours with rotation. For the final 30 minutes of incubation, Hoechst 33342 (Thermo, 62249) and BODIPY 493/503 (Invitrogen, D3922) were added at 1:500 dilutions. Adipocytes were washed twice and resuspended in 300 µL Fluoromount G (Southern Biotech, 0100-01) and mounted on glass slides with 1.4-1.6 mm concavity wells (Electron Microscopy Sciences, 71878-03). A sample of adipocytes was also incubated as above but without primary antibody to verify the specificity of the secondary antibody. Fluorescence images were acquired using Zeiss LSM 880 Upright Laser Scanning Confocal Microscope with filter cubes for DAPI, GFP, and Rhodamine in parallel using the 20X objective and processed using Zen Black 2.3 software. Images were analyzed and counted with ImageJ v. 1.53k.

### *Ex vivo* differentiation and transcriptional and high-content image-based characterization of differentiating primary human adipocyte progenitors

We obtained adipocyte progenitors from subcutaneous and visceral adipose tissue from patients undergoing a range of abdominal laparoscopic surgeries (sleeve gastrectomy, fundoplication or appendectomy). The visceral adipose tissue is derived from the proximity of the angle of His and subcutaneous adipose tissue obtained from beneath the skin at the site of surgical incision.

Additionally, human liposuction material was obtained. Each participant gave written informed consent before inclusion and the study protocol was approved by the ethics committee of the Technical University of Munich (Study № 5716/13). Isolation of AMSCs was performed as previously described^28^, and cells were differentiated in culture over 14 days. *Ex vivo* differentiated adipocytes were stained and imaged, and features were extracted using LipocyteProfiler as described in Laber et al. RNA-sequencing libraries were prepared and sequenced and QC’ed as previously described^28^. Bulk-RNA sequencing counts from subcutaneous and visceral samples differentiated for 14 days were deconvoluted using both subcutaneous and visceral adipocytes as reference as described above. Raw images collected during LipocyteProfiler analysis were randomly selected from samples predicted to have high or low content of hAd3, hAd5, or hAd6 adipocytes, and pseudocolored and combined using Adobe Photoshop.

### Gene Pathway Analysis

Analysis of enriched pathways in adipocyte markers was performed using clusterProfiler^68^ (version 3.16.1). Adipocyte cluster markers were filtered to an adjusted *p*-value < .05, then evaluated for enrichment in GO biological pathways or KEGG pathways containing under 300 genes.

### Identification and analysis of EBF2 SNP association with visceral adiposity

VAT, ASAT, and GFAT volumes in 40,032 individuals from the UK Biobank^69, 70^ who underwent MRI imaging were quantified as described elsewhere^71^. Variant rs4872393 was identified as a lead SNP associated with VATadjBMI and waist-to-hip ratio from summary statistics of two prior studies^31, 72^. Among the cohort who underwent MRI imaging, all variants at this locus (± 250 kb around rs4872393) with MAF >= 0.005 and imputation quality (INFO) score >= 0.3 were analyzed. For all 554 nominally significant (P < 0.05) variants associated with VATadjBMI in this region, a secondary conditional analysis testing for association with VATadjBMI was performed controlling for rs4872393 carrier status (P < 0.05/554 = 9 x 10^-5^).

Participants were excluded from analysis if they met any of the following criteria: (1) mismatch between self-reported sex and sex chromosome count, (2) sex chromosome aneuploidy, (3) genotyping call rate < 0.95, or (4) were outliers for heterozygosity. Up to 37,641 participants were available for analysis. Fat depot volumes adjusted for BMI and height (“adj” traits) were calculated by taking the residuals of the fat depot in sex-specific linear regressions against age at the time of MRI, age squared, BMI, and height^31^. Each trait was scaled to mean 0 and variance 1 in sex-specific groups before being combined for analysis. Linear regressions between a given trait-variant pair were adjusted for age at the time of imaging, age squared, sex, the first 10 principal components of genetic ancestry, genotyping array, and MRI imaging center. Analyses were performed using R 3.6.0 (R Project for Statistical Computing). *EBF2* regional visualization plot was made with the LocusZoom online tool^73^.

### Calculation of pseudobulk datasets to estimate adipose innervation

Approximate bulk RNA-seq datasets (pseudobulk) were obtained for visceral sNuc-seq samples by summing the total expression per-gene across all droplets containing a valid 10X cell barcode. This includes all cells that would normally have been removed in the single-nuclei studies by any of the filtering criteria (above): doublet score, splicing content, droplets with fewer than 400 UMIs, etc, in order to preserve the ambient RNA present in otherwise empty droplets. Repeated UMIs were still collapsed into single counts (per-droplet) before summing. Levels of pan-neuronal markers were calculated using this pesudobulk dataset and plotted against the proportion of visceral populations hAd2 and hAd6 relative to total adipocytes in each sample.

### Prediction of cell-cell interactions

Analysis of cell-cell interactions was performed using CellphoneDB^41^ (version 2.0.0). For human data, sNuc-seq counts data was split into files containing cells from subcutaneous and visceral fat from individuals with BMI lower than 30 or higher than 40. CellphoneDB with statistical analysis was run on each file separately to evaluate interactions in each condition. For mouse data, counts data was split into files containing cells from the inguinal and perigonadal fat of chow and high fat diet fed mice. Mouse gene names were converted to human gene names, as above, before running CellphoneDB with statistical analysis on each file.

### Identification of candidate etiologic cell types using CELLEX and CELLECT

CELLECT (https://github.com/perslab/CELLECT) and CELLEX (https://github.com/perslab/CELLEX) were used to identify candidate etiological cell types for a total of 23 traits. The input data for CELLECT is GWAS summary statistics for a given trait and cell type expression specificity (ES) estimates derived from single-cell RNA-seq data. The output is a list of prioritized candidate etiologic cell types for a given trait. ES estimates were calculated using CELLEX (version 1.1), which computes robust estimates of ES by relying on multiple expression specificity measures (for further details see Timshel et. al.^74^). CELLEX was run separately on the raw mouse and human (sNuc) gene expression matricies to compute gene expression specificities for each cluster based on the clustering assignment reported above. The resulting cell type specificity matrix was used along with multiple GWAS studies^30, 75–79^ (**Extended Data Table 3**) as input for CELLECT^74^ (version 1.1), which was run with default parameters. Significant cell types were identified using a by-trait and by-species Bonferroni *p*-value threshold of *p*<0.05.

### SNP analysis for bulk mRNA-seq cohort

The raw GTC SNP expression data from Infinium OmniExpress-24 Kit was converted to VCF format using Picard version 2.21.6. The pre-processing of the SNP data before phasing and imputation was performed using plink2 (https://www.cog-genomics.org/plink/2.0/). The SNP genotype was then phased and imputed using the Eagle v2.3.5^80^ and Minimac3^81^ packages, respectively. SNPs were mapped to the NCBI database using the rsnps package (https://CRAN.R-project.org/package=rsnps) and filtered to keep only SNPs that had a minor allele frequency > 0.05. For plotting gene expression against genotype, bulk RNA sequencing data was TMM normalized using edgeR^82^. Statistical validation for significance was done using the Wilcoxon rank-sum Test which is a non-parametric test assuming independent samples.

### Statistics

*p*-values for scatterplots were calculated using GraphPad Prism version 8.0 and represent the probability that the slope of the line of best fit is nonzero. All error bars on bar graphs represent standard error. Statistics on proportional composition graphs were calculated using scCODA^83^ (version 0.1.2) using the Hamiltonian Monte Carlo sampling method. The model formula used was “Depot + BMI” (human) or “Depot + Diet) (mouse) for all objects in for which both of these covariates were present, or the individual covariate when only a single condition was present.

## DATA AVAILABILITY

Single cell RNA expression and count data is deposited in the Single Cell Portal (Study #<u>SCP1376</u>) and will be downlodable upon publication. Processed count data for bulk RNA-seq and dge matrices for single cell and single nucleus RNA-seq have been deposited in GEO and will be made public upon publication (Bulk-seq Accession #GSE174475, sc-RNA-seq/sNuc-sec Acession #GSE176171), raw sequencing reads for mouse data will additionally be deposited before publication. FASTQ and SNP array files for human samples will be deposited in dbGaP before publication.

## CODE AVAILABILITY

Data analysis pipelines used in this study for processing of raw sequencing data, integration, and clustering can be obtained from https://gitlab.com/rosen-lab/white-adipose-atlas.

## ACKNOWLEDGEMENTS

This work was supported by NIH grants RC2 DK116691 to EDR, LTT, AC, OA, and AR, AHA POST14540015 and DoD PRMRP-DAW81XWH to LTT, Broad-BADERC Collaboration Initiative Award (NIH 5P30DK057521) to LTT and EDR, and R01 DK102173 to EDR. MPE is supported by NIH grant F32DK124914. Additional support includes PRIN 2017 (Italian Ministry of University, #2017L8Z2EM) to AG, THP acknowledges the Novo Nordisk Foundation (unconditional donation to the Novo Nordisk Foundation Center for Basic Metabolic Research; grant number NNF18CC0034900) and the Lundbeck Foundation (Grant number R190-2014-3904), grants AMP-T2D RFB8b (FNIH) and UM1DK126185 (NIDDK) to MC, Sarnoff Cardiovascular Research Foundation Fellowship to S.A., grants 1K08HG010155 and 1U01HG011719 to A.V.K. from the National Human Genome Research Institute, and a sponsored research agreement from IBM Research to the Broad Institute of MIT and Harvard to A.V.K. All single cell library construction and sequencing was performed through the Boston Nutrition Obesity Research Center Functional Genomics and Bioinformatics Core (NIH P30DK046200). We thank Christina Usher for artistic support and Miriam Udler for helpful discussions.

## AUTHOR CONTRIBUTIONS

MPE, LTT, and EDR conceived of the project. MPE and EDR wrote the manuscript with assistance from LTT, CJ, OA, and AR. MPE, ALE, DP, DT, GC, ADV, AS, EM, SS, SL, GPW, MLV, and AGu performed experiments. GPW, AGu, ZK, JD, CGB, WG, AC, SJL, BTL, DM, and AT collected samples. MPE, CJ, AMJ, HD, SA, AK, and HS performed computational analysis. AVK, MC, THP, AGi, OA, and AR provided additional intellectual input.

## ADDITIONAL INFORMATION

Supplementary information is available for this paper.

Correspondence and requests for materials should be addressed to EDR.

## EXTENDED DATA FIGURE AND TABLE LEGENDS

**Extended Data Fig. 1.**
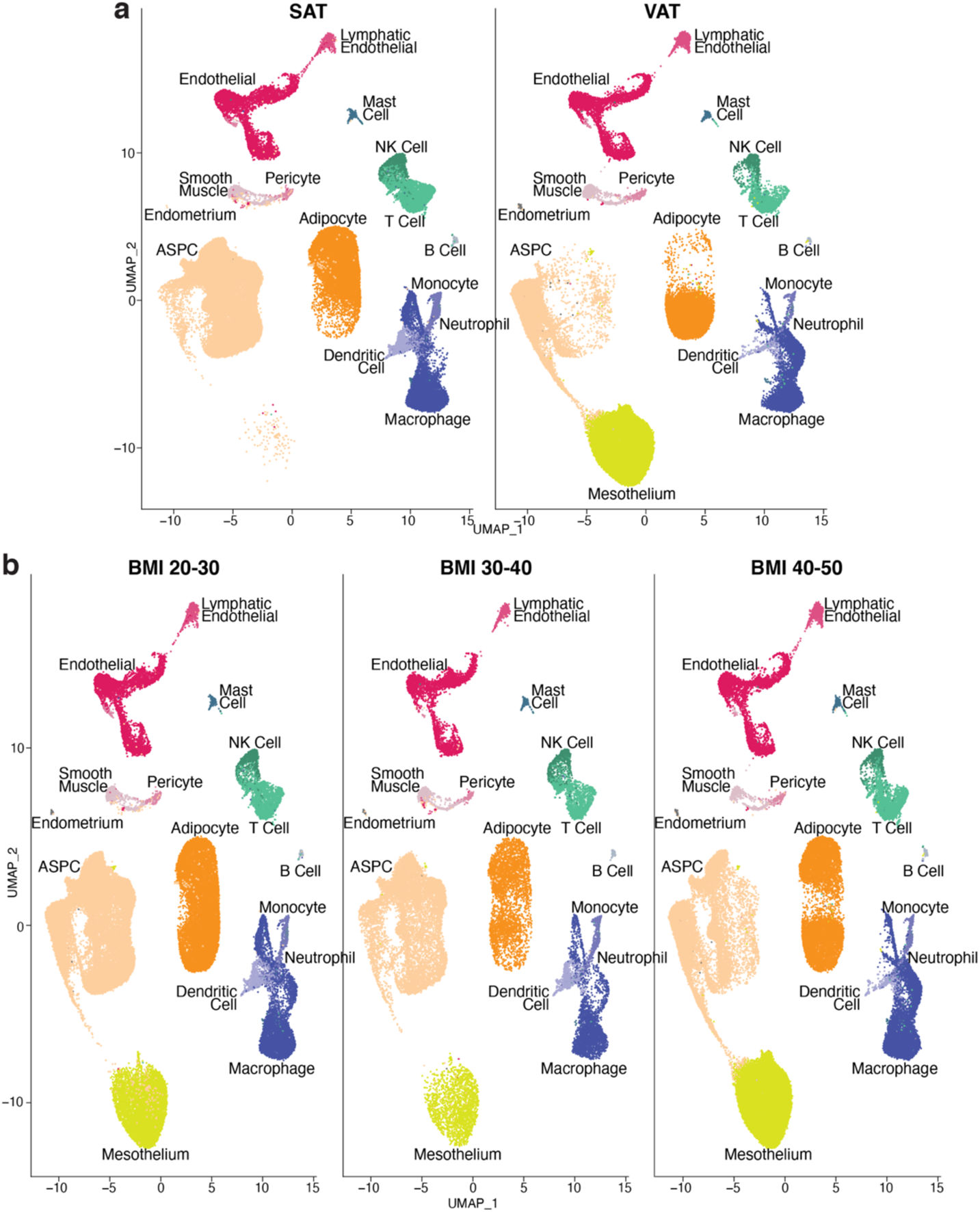
Recovery of human WAT cell types is highly influenced by adipose depot. **a,** UMAP projection of all human cells split by depot. **b,** UMAP projection of all human cells split by BMI range.

**Extended Data Fig. 2.**
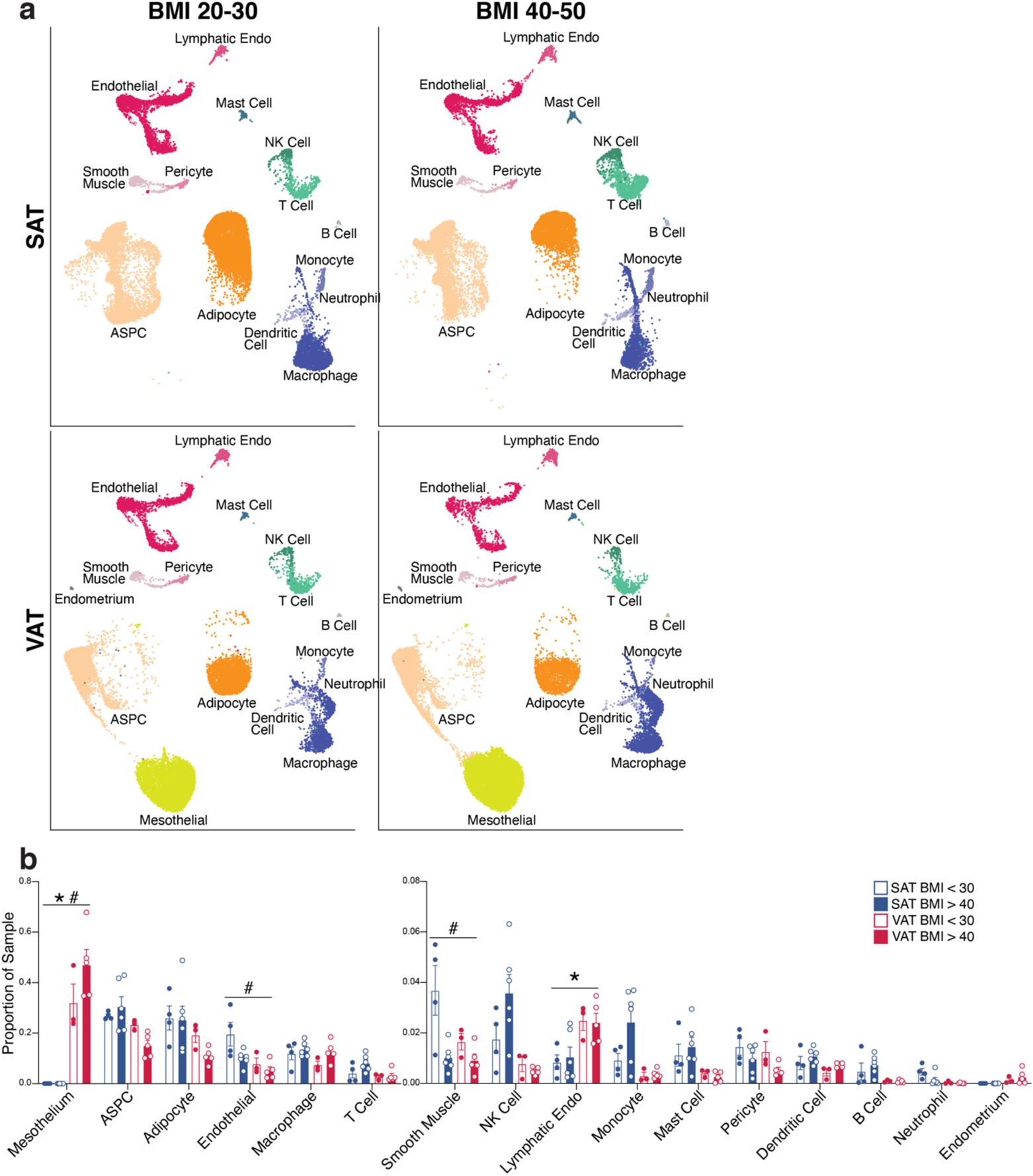
Additional analysis of the effects of depot and BMI on human WAT populations. **a,** UMAP projections of cells from the lowest and highest BMI ranges in the dataset, split by depot. To facilitate comparison, samples were randomly subset to contain the same number of cells in each plot (n = 20,339). **b,** Graph showing the proportion of sNuc-seq cells in each cluster per sample, split by depot and BMI. For bar graphs, * indicates credible depot effect and # indicates credible BMI effect, calculated using dendritic cells as reference.

**Extended Data Fig. 3.**
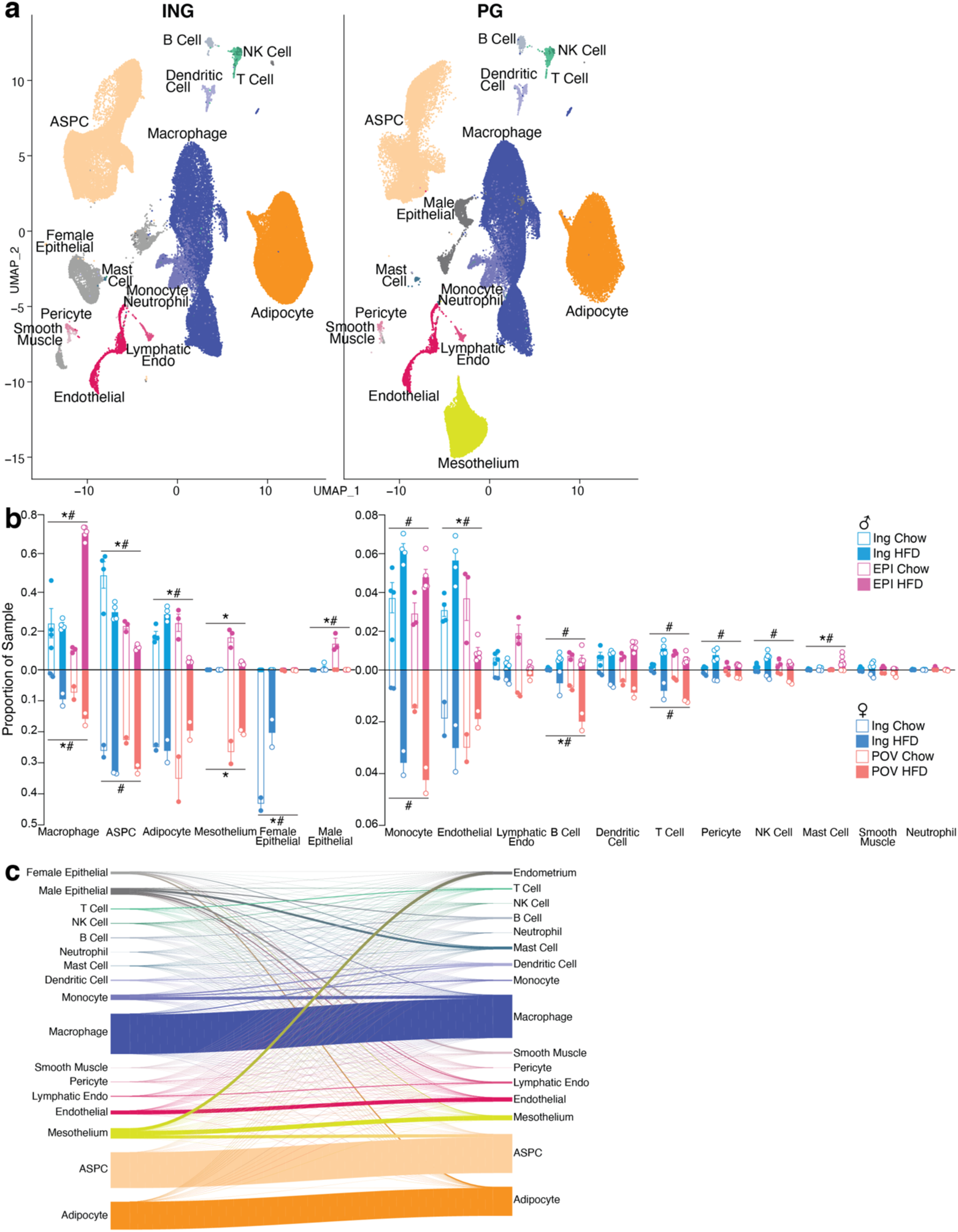
Additional analysis of the effects of depot and diet on mouse WAT populations and association with human WAT populations. **a,** UMAP projection of all mouse WAT cells split by depot. **b,** Proportion of cells in each cluster per sample, split by sex as well as by depot and diet. **c,** Riverplot showing the relationship between mouse and human clusters. Mouse cells were mapped onto human sNuc-seq cells using multimodal reference mapping. The riverplot represents the relationship between manually assigned mouse cluster and mapped human cluster for every mouse cell. For bar graph, error bars represent SEM, * indicates credible depot effect and # indicates credible diet effect, calculated using dendritic cells as reference.

**Extended Data Fig. 4.**
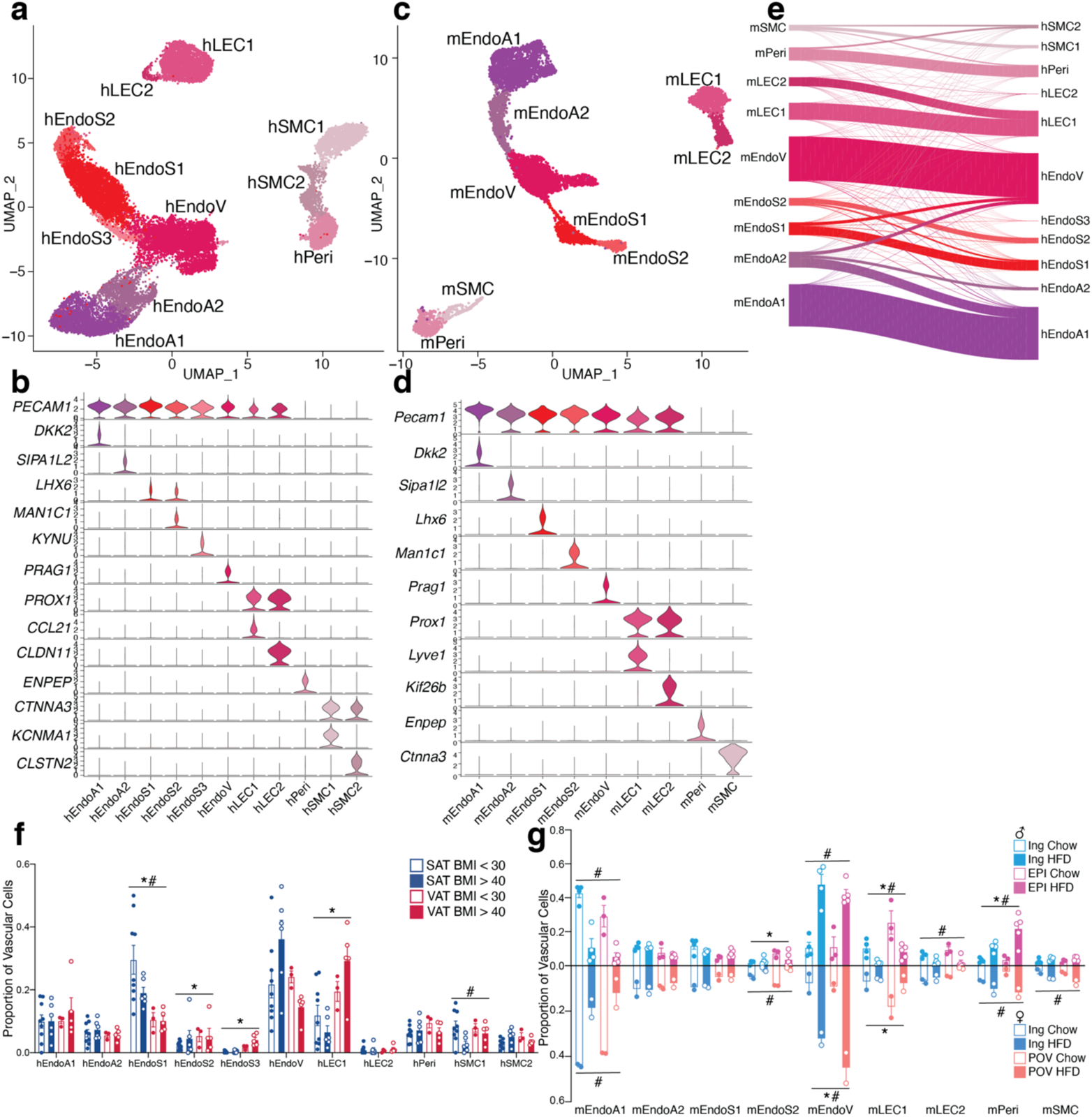
Highly similar vascular cells in human and mouse WAT. **a,** UMAP projection of 22,734 human vascular cells. **b**, Marker genes for 11 distinct clusters of human WAT vascular cells. **c,** UMAP projection of 7,632 mouse vascular cells. **d, M**arker genes for 9 distinct clusters of mouse WAT vascular cells. **e,** Riverplot showing the correlation between annotated mouse and human vascular clusters based on multimodal reference mapping for each mouse cell. **f-g**, Bar graphs showing the proportion of cells in each cluster per sample split by depot and BMI for human (**f**) and depot, diet, and sex for mouse (**g**). For bar graphs, error bars represent SEM, * indicates credible depot effect and # indicates credible BMI/diet effect, calculated using hEndoA2 (human) and mEndoA2 (mouse) as reference.

**Extended Data Fig. 5.**
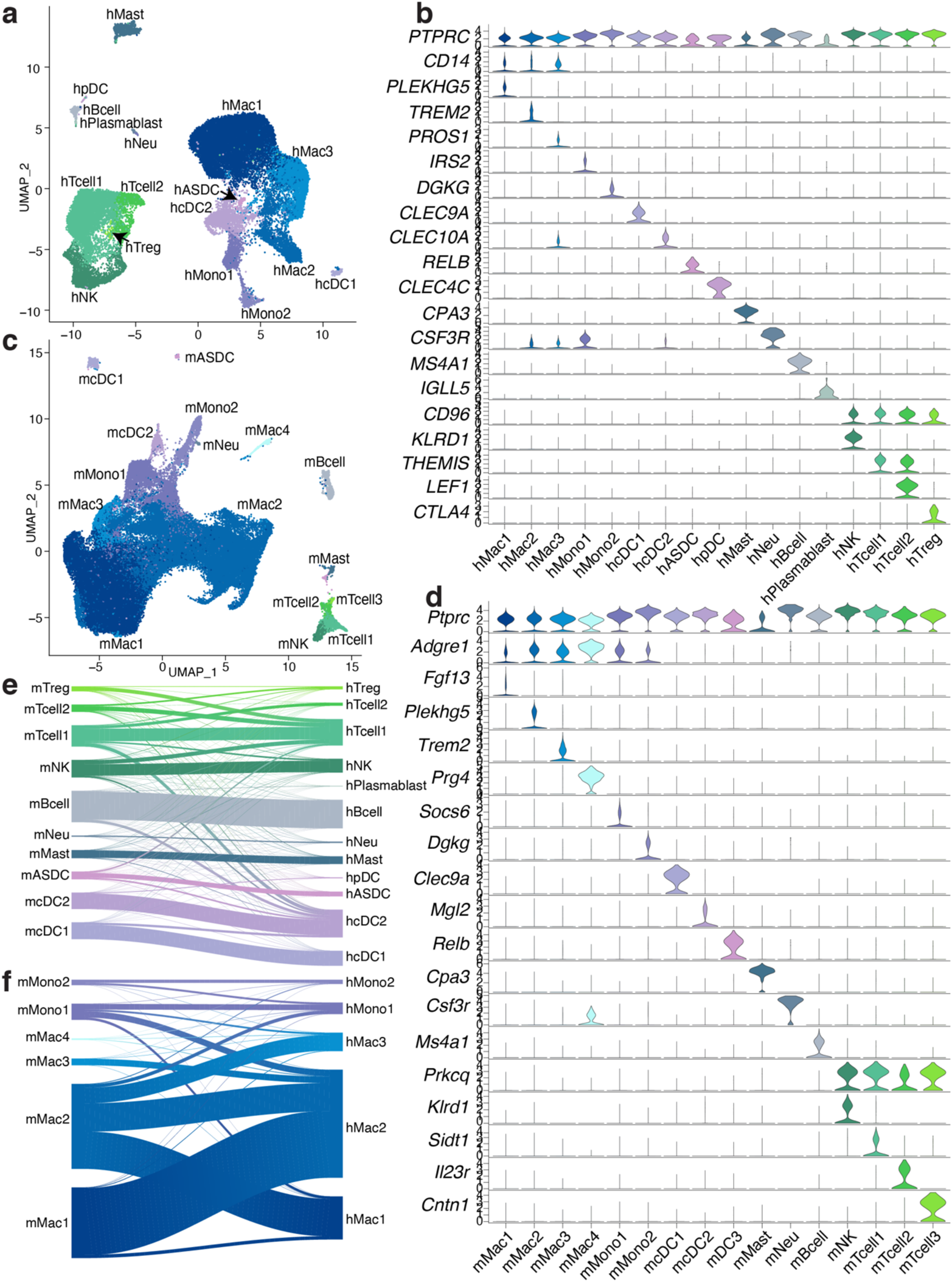
Comparison of immune cells in human and mouse WAT. **a,** UMAP projection of 34,268 immune cells from human WAT. **b,** Marker genes for human immune cell clusters. **c,** UMAP projection of 70,547 immune cells from mouse WAT. **d,** Marker genes for mouse immune cell clusters. **e-f,** Riverplots showing the correlation between annotated mouse cluster and mapped human cluster for mouse (**e**) dendritic cells, mast cells, neutrophils, B cells, NK cells, and T cells and (**f**) monocytes and macrophages.

**Extended Data Fig. 6.**
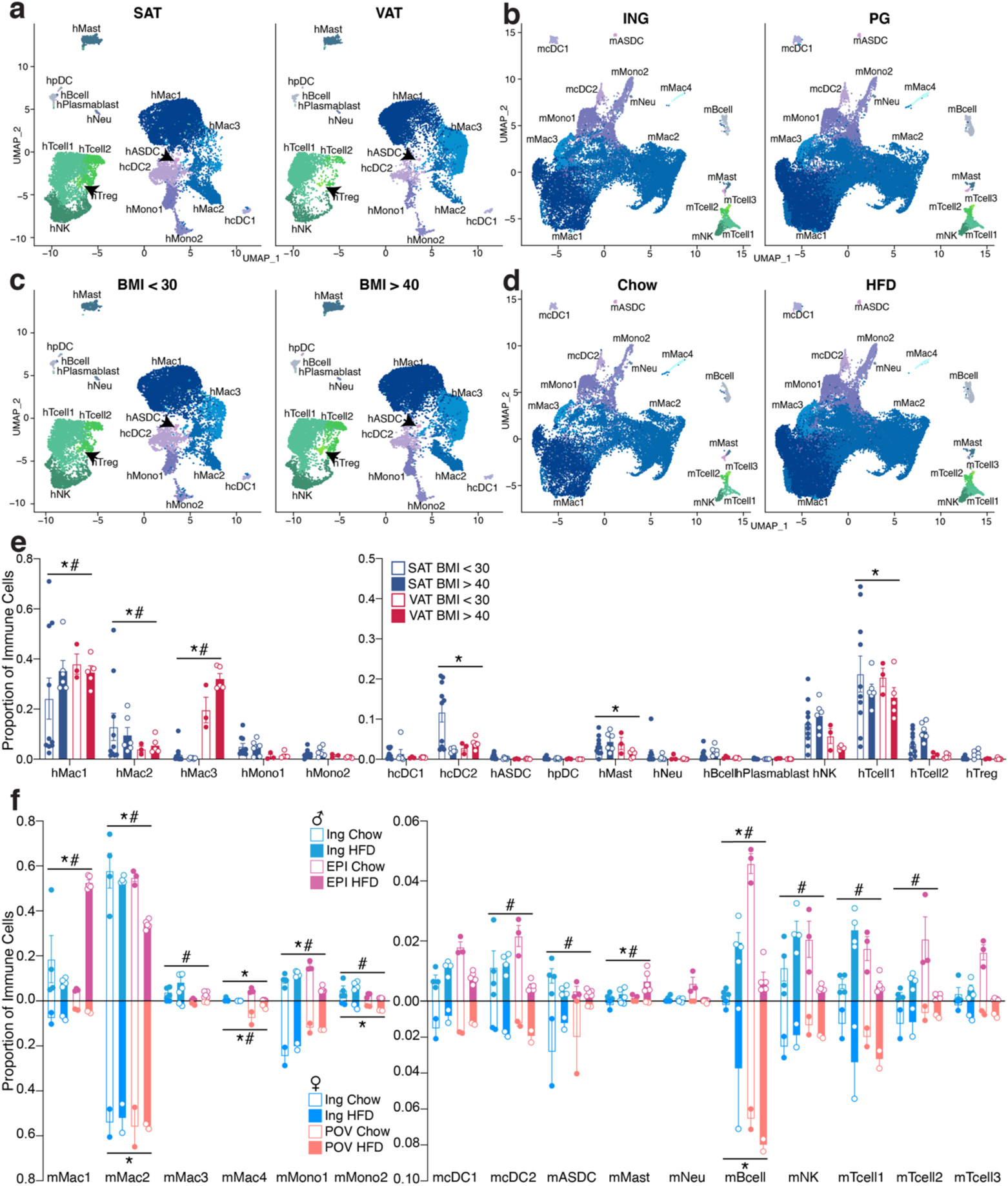
Human and mouse immune cells are differentially regulated by depot and BMI/diet. **a-b,** UMAP projections of human (**a**) and mouse (**b**) WAT immune cells split by depot. **c-d,** UMAP projections of human (**c**) and mouse (**d**) WAT immune cells split by BMI (**c**) and diet (**d**). **e-f,** Bar graphs showing the proportion of cells in each cluster per sample split by depot and BMI for human (**e**) and depot, diet, and sex for mouse (**f**). For bar graphs, error bars represent SEM, * indicates credible depot effect and # indicates credible BMI/diet effect, calculated using hMono2 (human) and mcDC1 (mouse) as reference.

**Extended Data Fig. 7.**
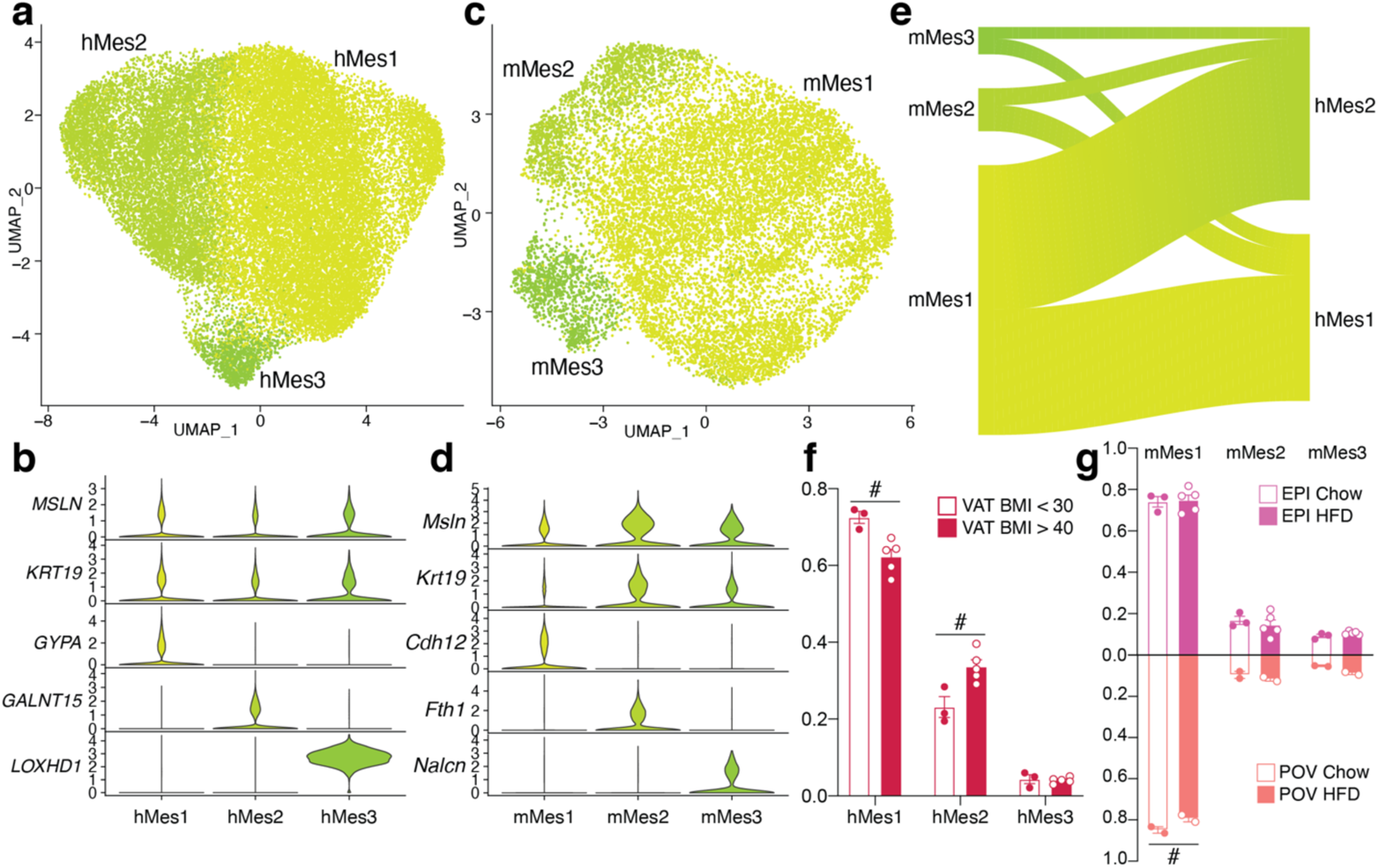
Subpopulations of human and mouse mesothelial cells. **a,** UMAP projection of 30,482 human mesothelial cells. **b,** Marker genes for distinct human mesothelial populations. **c,** UMAP projection of 14,947 mouse mesothelial cells. **d** Marker genes for distinct mouse mesothelial populations. **e,** Riverplots showing relationship of mouse and human mesothelial clusters. **f-g,** Proportion of cells in each cluster per sample, split by BMI for human (**f**) and diet and sex for mouse (**g**). Error bars represent SEM, # indicates credible BMI/diet effect, calculated using hMes3 (human) and mMes1 (mouse) as reference.

**Extended Data Fig. 8.**
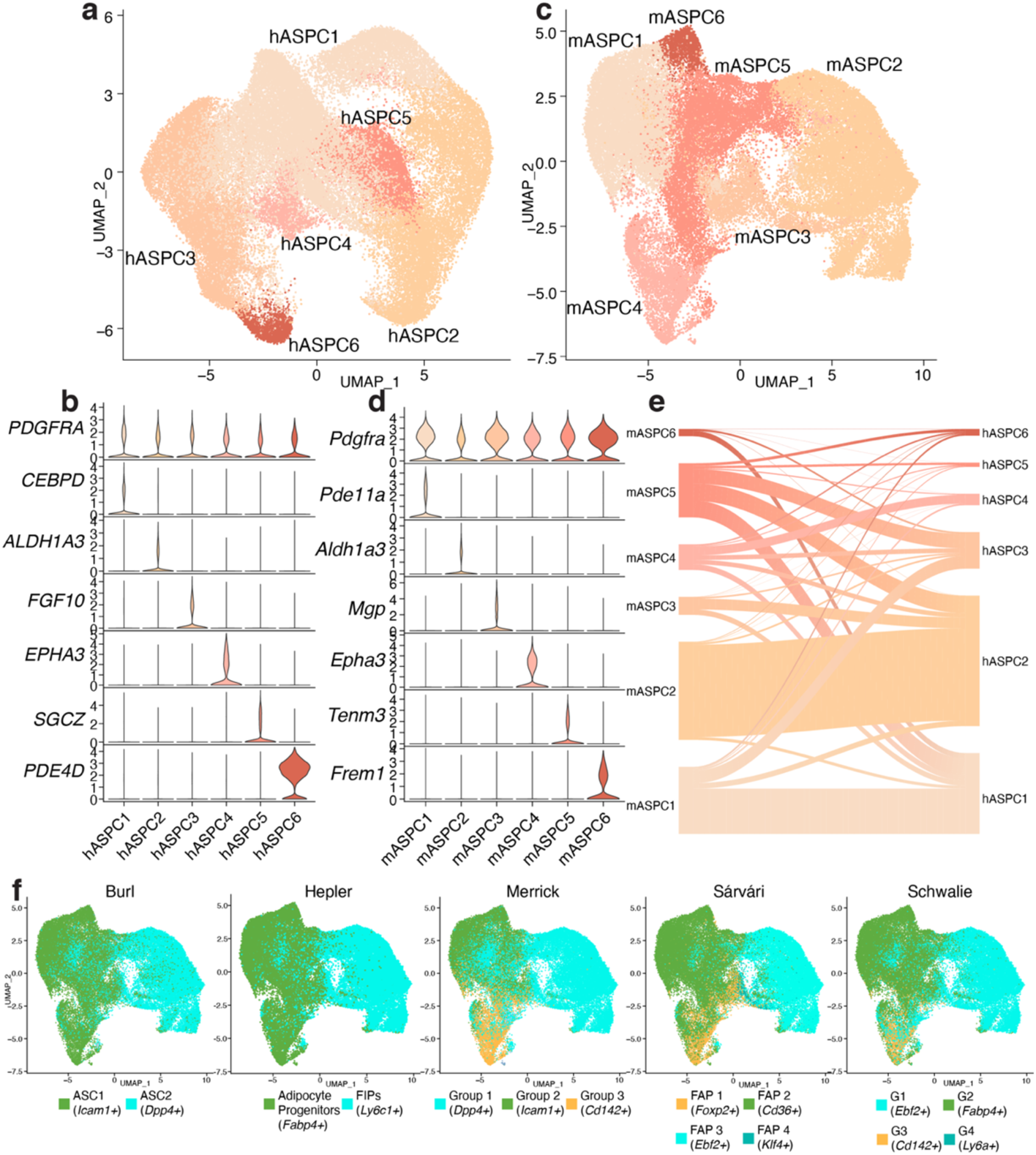
Human and mouse ASPCs share commonalities with previously reported subtypes. **a,** UMAP projection of 52,482 human ASPCs. **b**, Marker genes for distinct ASPC subpopulations. **c,** UMAP projection of 51,227 mouse ASPCs. **d**, Marker genes for distinct ASPC subpopulations. **e,** Riverplot depicting the relationship between mouse and human ASPC clusters. **f,** Reference mapping of ASPC cell types reported by other groups onto the mouse ASPCs from this paper.

**Extended Data Fig. 9.**
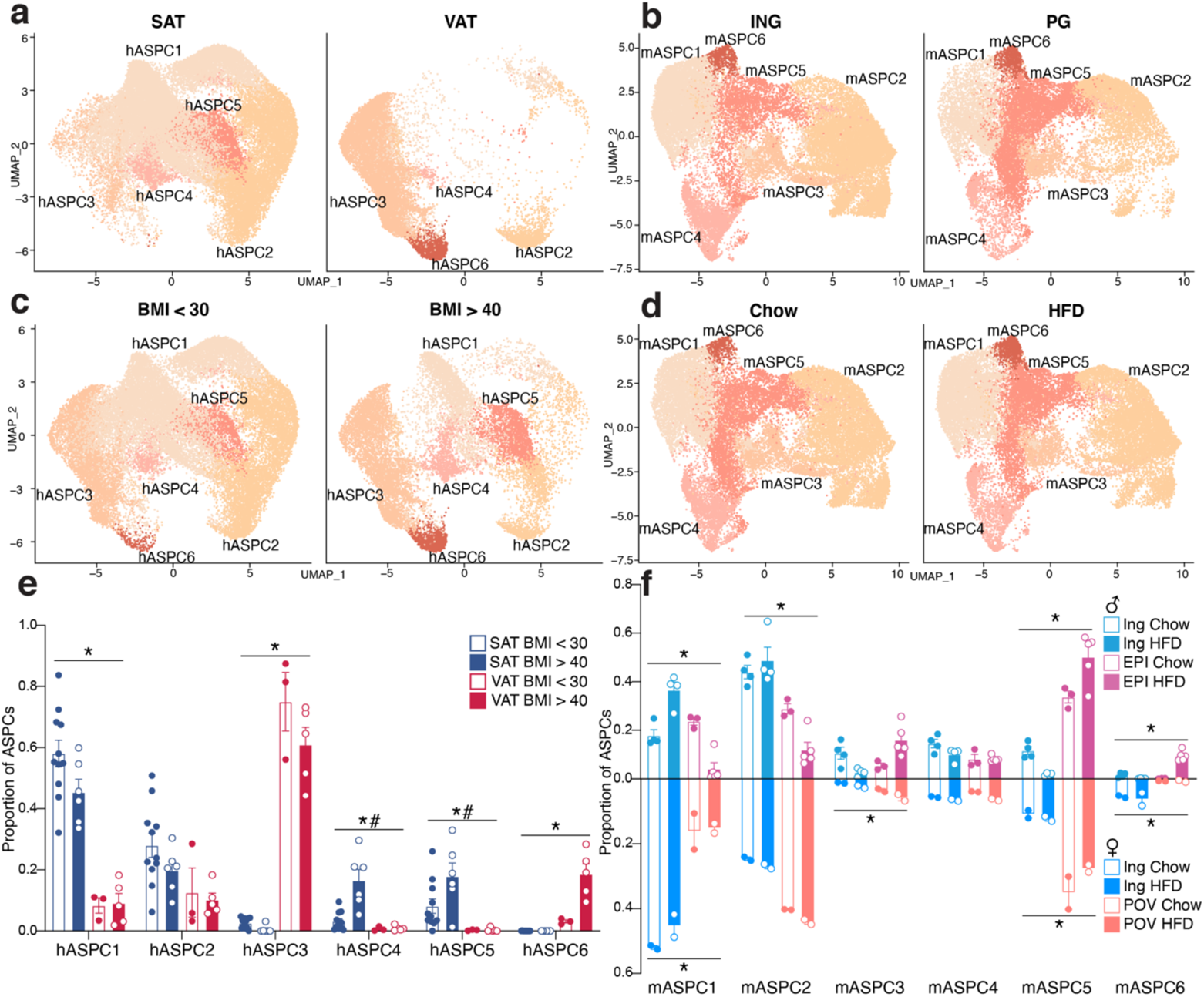
Human ASPCs exhibit strong depot dependency while mouse ASPCs are dependent on both depot and diet. **a-b,** UMAP projections of human (**a**) and mouse (**b**) ASPCs split by depot. **c-d,** UMAP projections of human (**c**) and mouse (**d**) ASPCs split by BMI/diet. **e-f,** Proportion of ASPC cells in each cluster per sample split by depot and BMI for human (**e**) and depot, diet, and sex for mouse (**f**). For bar graphs, error bars represent SEM, * indicates credible depot effect and # indicates credible BMI/diet effect, calculated using hASPC2 (human) and mASPC4 (mouse) as reference.

**Extended Data Fig. 10.**
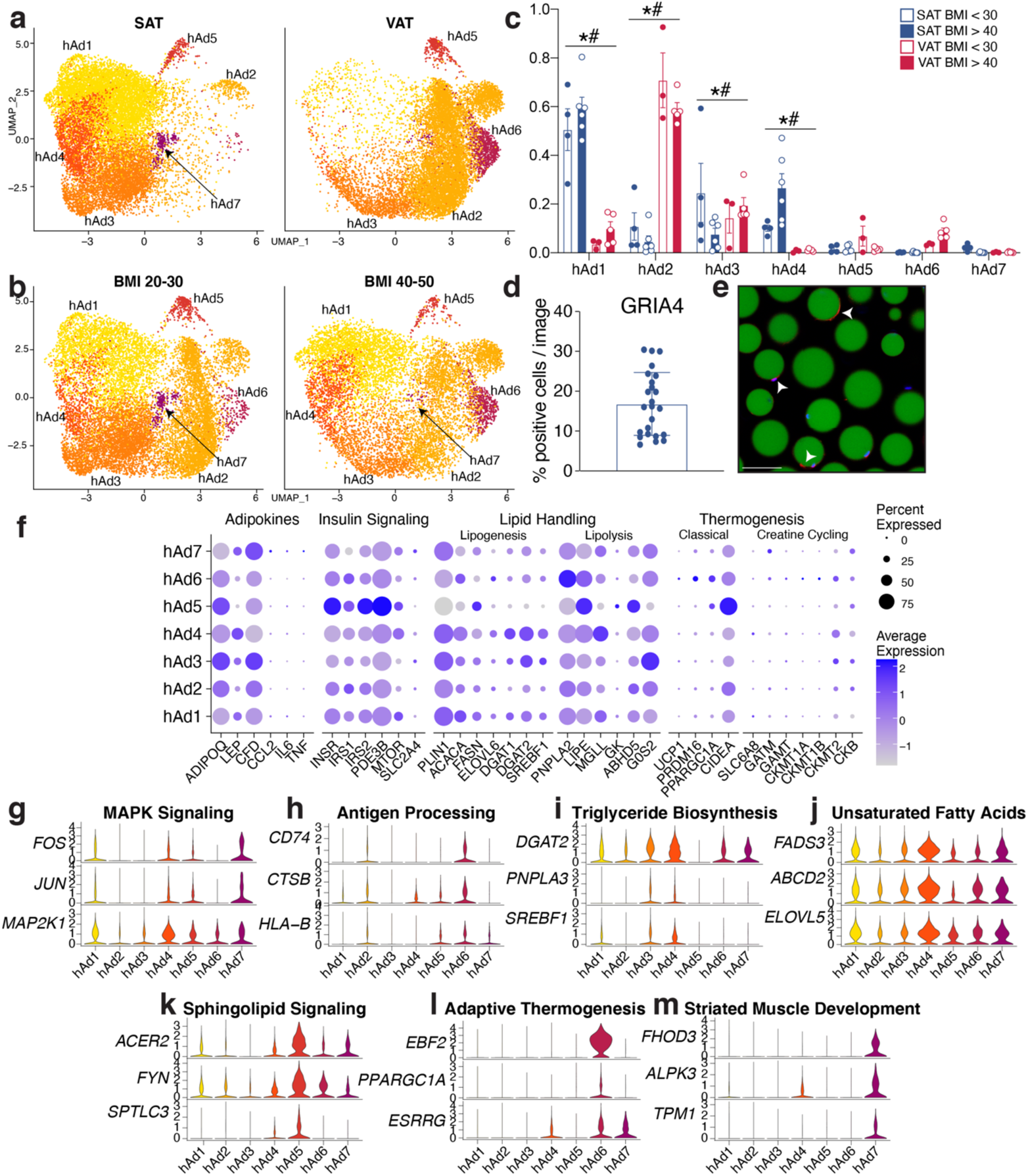
Human adipocyte subtypes are highly dependent on depot and may be responsible for distinct functions. **a-b,** UMAP projections of human white adipocytes split by depot (**a**) and BMI (**b**). **c,** Proportion of cells in each human cluster by sample split by depot and BMI. **d,** Quantification of immunofluorescence analysis of GRIA4+ cells in mature human adipocytes from two individuals. Each dot represents an image. **e,** Representative images of GRIA4+ cells. **f,** Expression of genes associated with adipokine secretion, insulin signaling, lipid handling, and thermogenesis across human adipocyte subclusters. **g-m,** Expression of genes associated with GO or KEGG pathways indicative of individual human adipocyte subclusters. For bar graph, error bars represent SEM, * indicates credible depot effect and # indicates credible BMI effect, calculated using hAd5 as reference.

**Extended Data Fig. 11.**
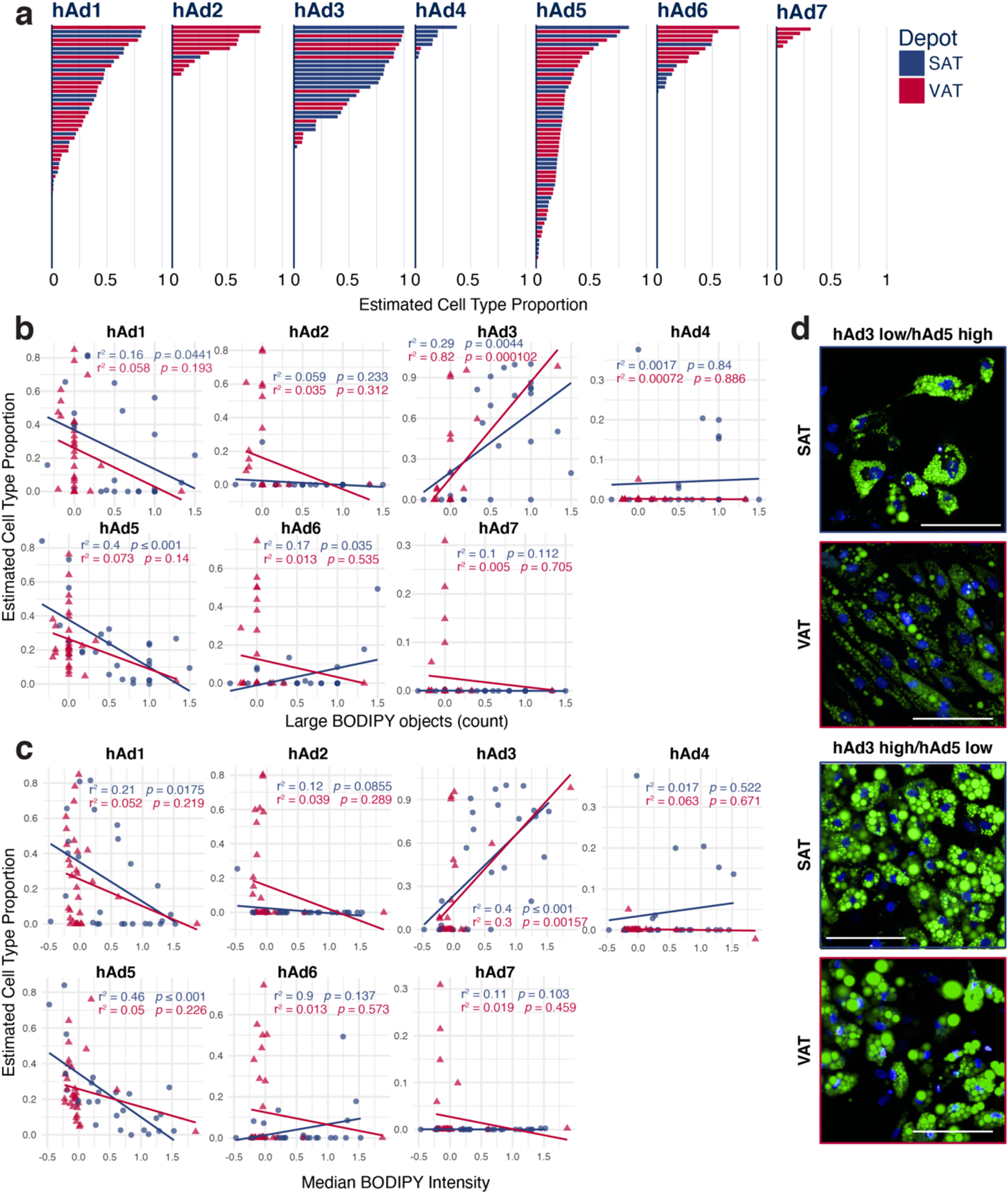
Human adipocytes differentiated ex vivo recapitulate many of the adipocyte subclusters found in vivo. **a**, Plot of estimated cell type proportion in ex vivo adipocyte cultures differentiated from subcutaneous or visceral preadipocytes for 14 days, ordered by estimated proportion. **b-c,** Scatterplots showing the relationship between estimated cell type proportion and the LipocyteProfiler-calculated features Large BODIPY objects (**b**) and Median BODIPY Intensity (**c**). **d,** Representative images of hAd3 low/hAd5 or hAd3 high hAd5 low in vitro differentiated cultures. Green represents BODIPY staining, blue represents Hoechst staining.

**Extended Data Fig. 12.**
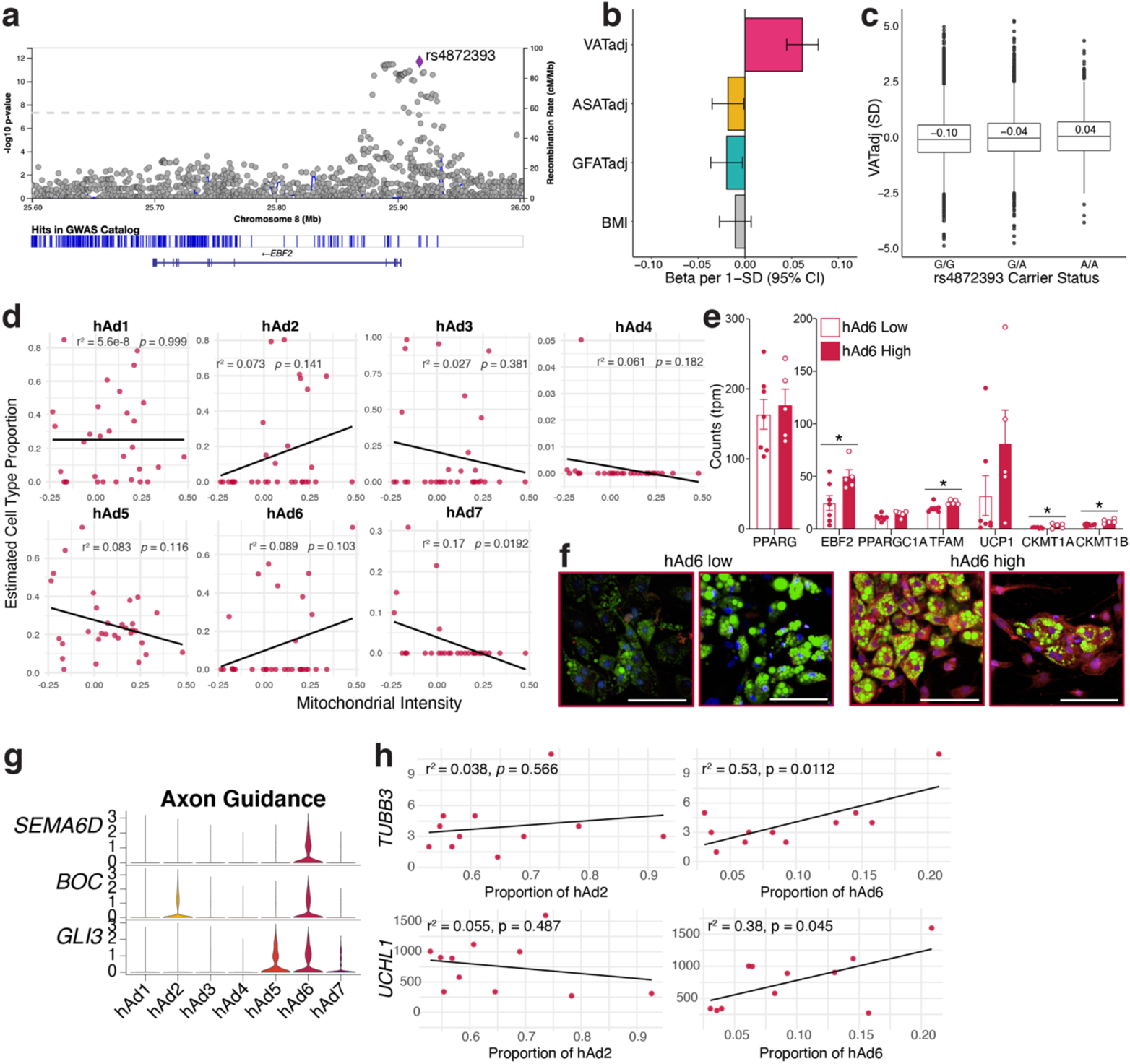
Visceral-specific adipocyte subpopulation hAd6 is associated with thermogenic traits. **a,** Regional visualization of associations of common genetic variants near EBF2 with VATadj. **b,** Association of rs4872393 with VATadj, ASATadj, GFATadj, and BMI per minor allele A; n = 37,641. **c,** VATadj raw data plotted according to rs4872393 carrier status; n = 36,185. **d,** Scatterplot showing the relationship between estimated cell type proportion and the LipocyteProfiler calculated feature Mitochondrial Intensity in visceral samples. **e,** Expression of mitochondrial and thermogenic genes in visceral in vitro differentiated adipocytes stratified by estimated hAd6 proportion and matched for amount of differentiation using *PPARG* levels. **f,** Representative images of hAd6 low and high visceral in vitro differentiated cultures. Green represents BODIPY staining, red represents MitoTracker staining, and blue represents Hoechst staining. **g,** Violin plot of sNuc-seq data showing axon guidance genes in adipocyte subclusters. **h,** Scatterplots showing the relationship between calculated proportion of visceral subpopulations hAd2 and hAd6 and expression of pan-neuronal markers on the ambient RNA of individual visceral sNuc-seq samples. For bar graph, error bars represent SEM, *, *p* < .05, **, *p* < .01.

**Extended Data Fig. 13.**
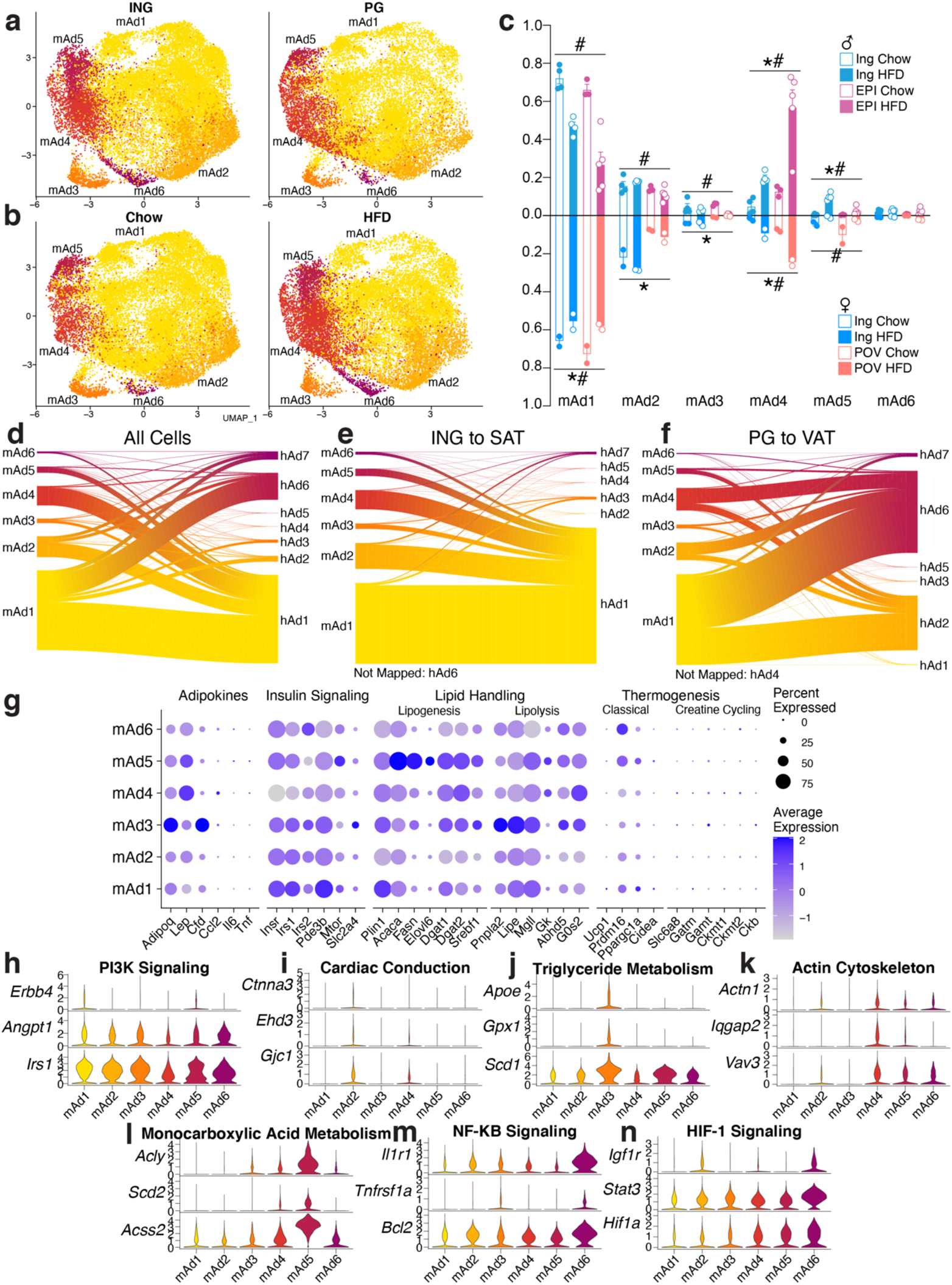
Mouse adipocytes appear to have distinct functionality but are not analogous to human adipocyte subpopulations. **a-b,** UMAP projections of mouse adipocytes split by depot (**a**) and diet (**b**). **c,** Proportion of cells in each mouse cluster per sample split by depot, diet, and sex. **d,** Expression of genes associated with known adipocyte functions in mouse adipocyte subclusters. **e-k,** Expression of genes associated with GO or KEGG pathways indicative of individual mouse adipocyte subclusters. **l-n,** Riverplots of mouse cells showing the association between mouse and human adipocyte clusters from both subcutaneous and visceral depots (**l**), subcutaneous (ING and SAT) adipocytes only (**m**) or visceral (PG and VAT) adipocytes only (**n**). For depot comparisons, both mouse query objects and human reference objects were subset to the respective depot before mapping. For bar graph, error bars represent SEM, * indicates credible depot effect and # indicates credible diet effect, calculated using mAd6 as reference.

**Extended Data Fig. 14.**
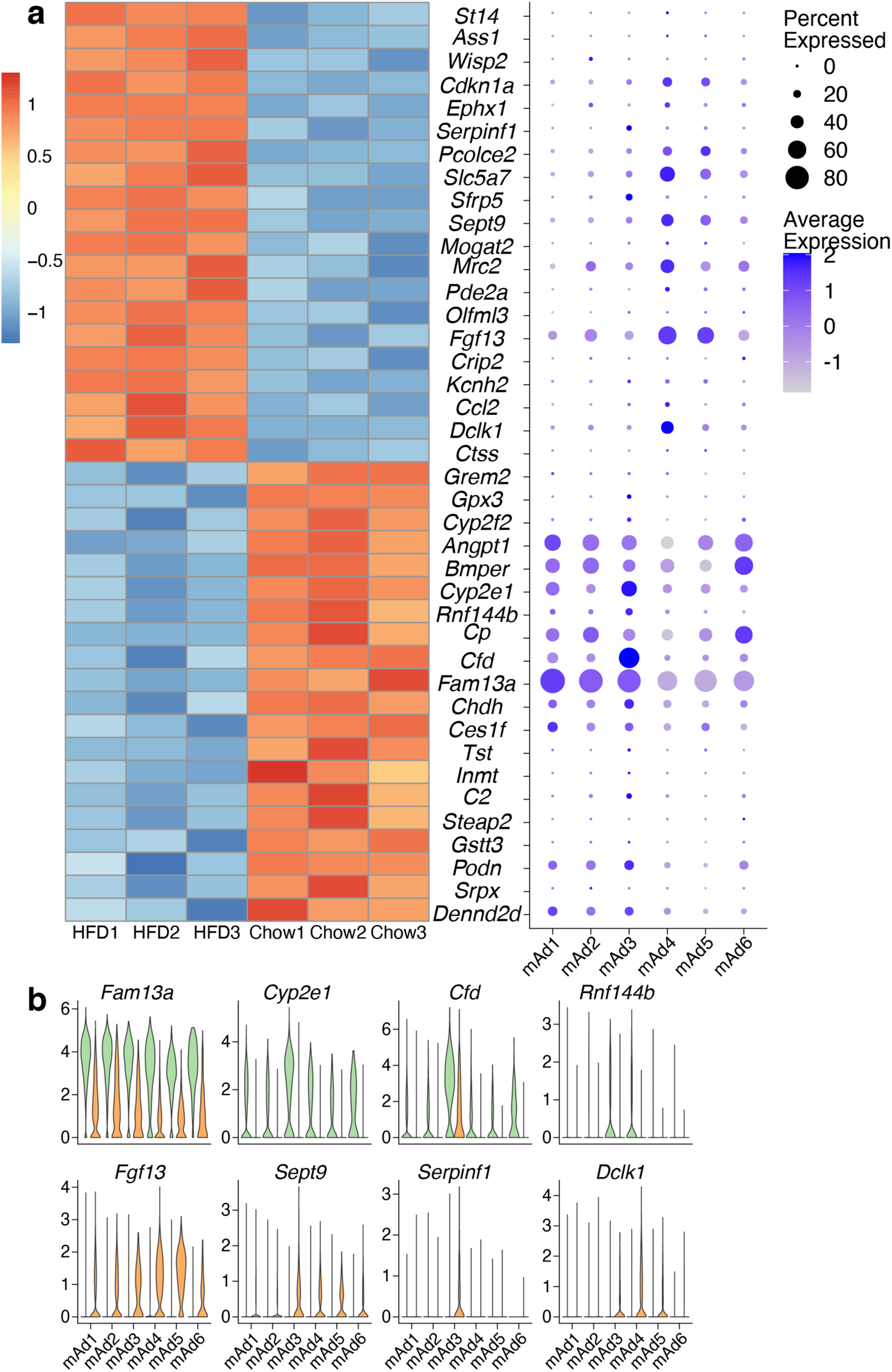
Adipocyte gene expression changes during high fat diet result from both changes in abundance of adipocyte subtypes and from expression changes within subclusters. **a,** (left) Heatmap depicting expression of the top 20 most up- and down-regulated genes in adipocytes after HFD feeding, as determined by bulk sequencing of TRAP-isolated adipocyte RNA. On the right, the same genes are plotted onto the mouse adipocyte subclusters to determine cluster specificity. **b,** Selected genes from **a** are plotted onto mouse adipocyte subclusters and split by diet.

**Extended Data Fig. 15.**
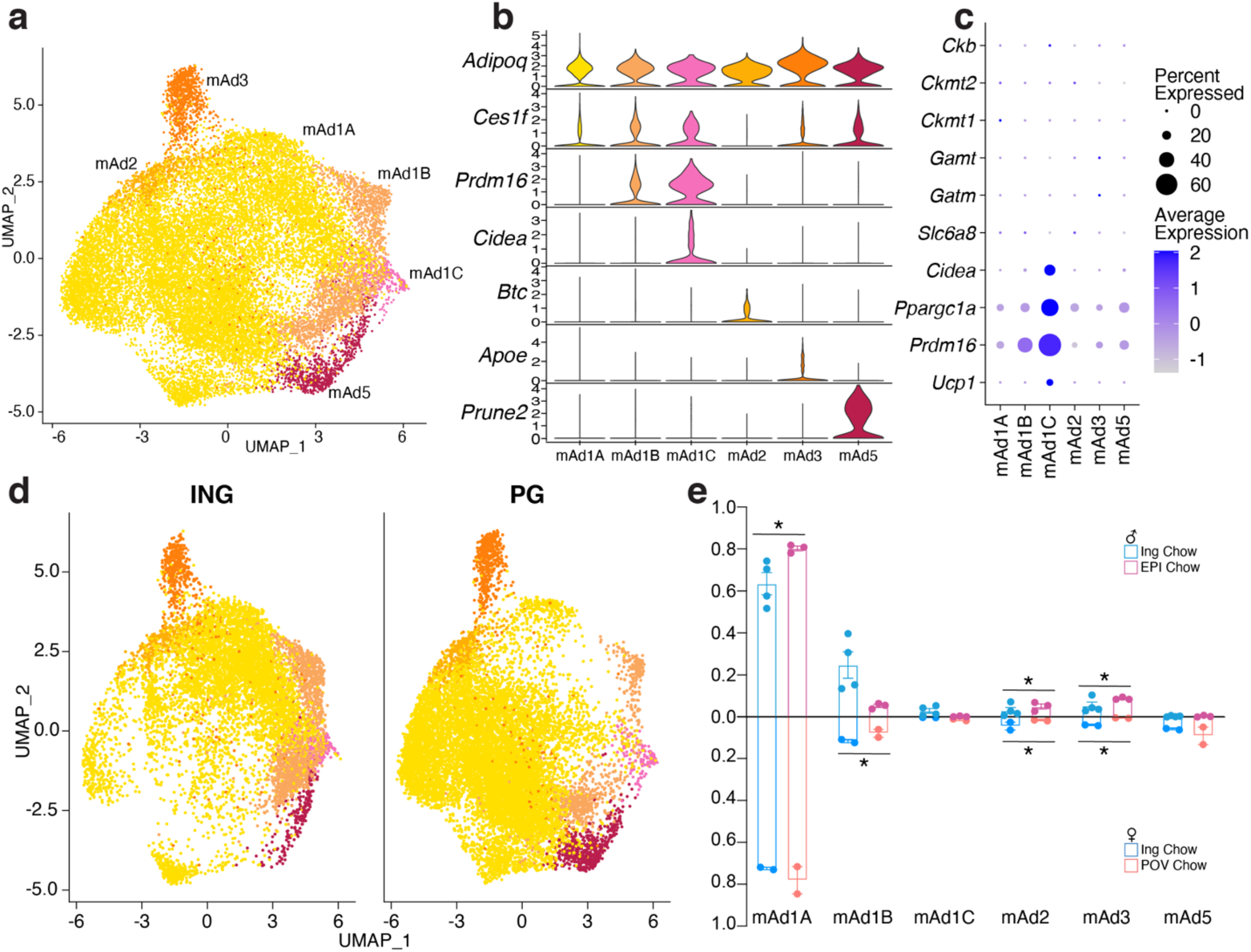
Mouse adipocytes from chow fed animals form a thermogenic subpopulation. **a,** UMAP projection of 21,519 adipocytes from chow fed animals. **b,** Marker gene expression of adipocytes from chow fed mice. **c,** Thermogenic gene expression in mouse chow adipocyte subclusters. **d,** UMAP projection of adipocytes from chow fed animals split by depot. **e,** Proportion of cells in each sample by cluster split by depot and sex. Error bars represent SEM, * indicates credible depot effect, calculated using mAd5 as reference.

**Extended Data Fig. 16.**
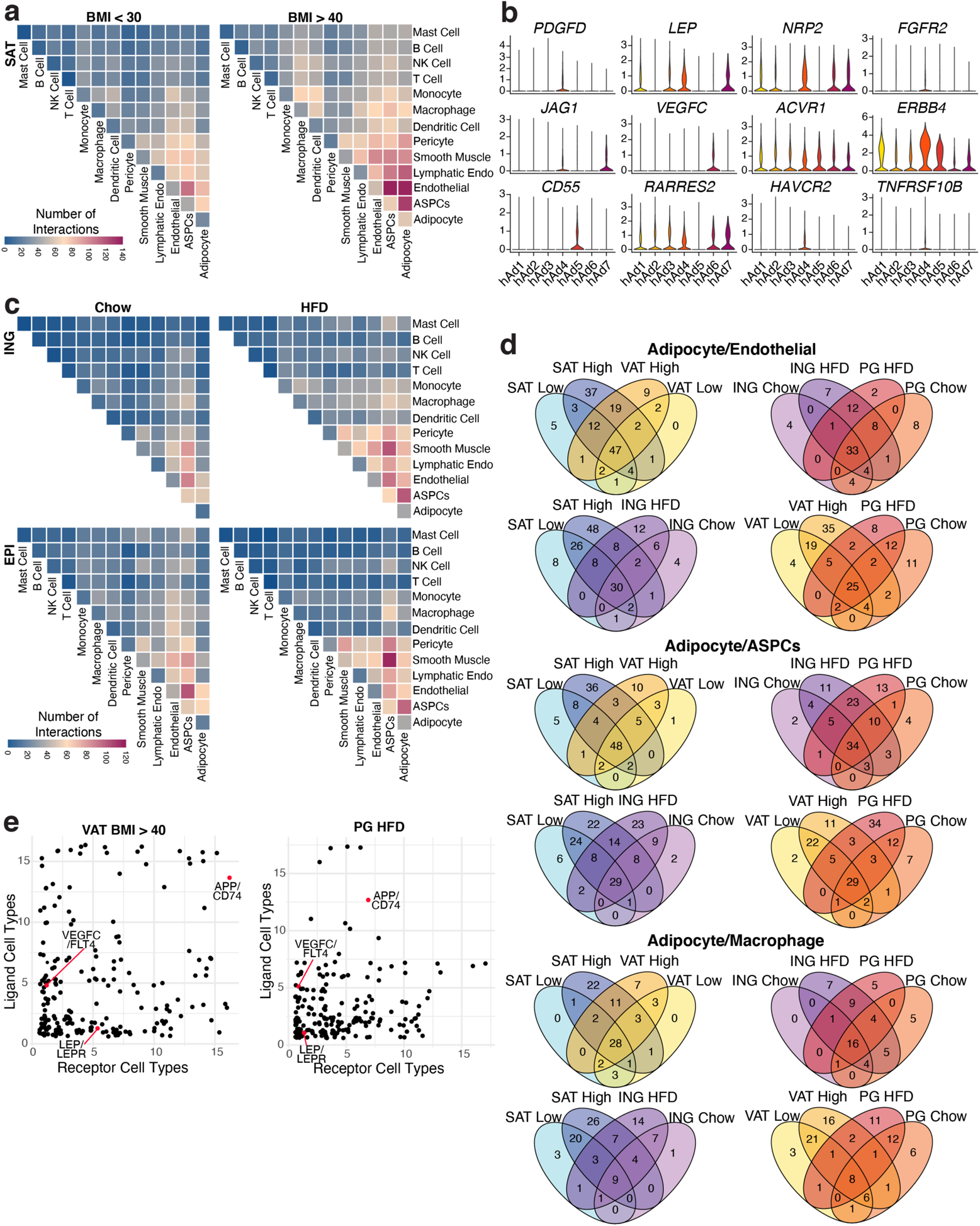
CellphoneDB identifies increasing numbers of cell-cell interactions within WAT during obesity. **a,** Heatmap showing number of significant interactions identified between cell types in SAT of low (<30) and high (>40) BMI individuals as determined by CellphoneDB. **b,** Expression levels of ligand and receptor genes from Figure 4b in human adipocyte subclusters. **c,** Heatmaps showing number of significant interactions identified between cell types in ING and PG WAT of chow and HFD fed mice. **d,** Venn diagrams showing the overlap of significant interactions between adipocytes and endothelial cells, ASPCs, and macrophages between depot, BMI/diet, and species. **e,** Jitter plots of the relationship between number of WAT cell types expressing a ligand (y axis) vs. the number of cell types expressing the receptor (x axis) for all significant interactions in high BMI human VAT (left) and mouse HFD PG (right).

**Extended Data Fig. 17.**
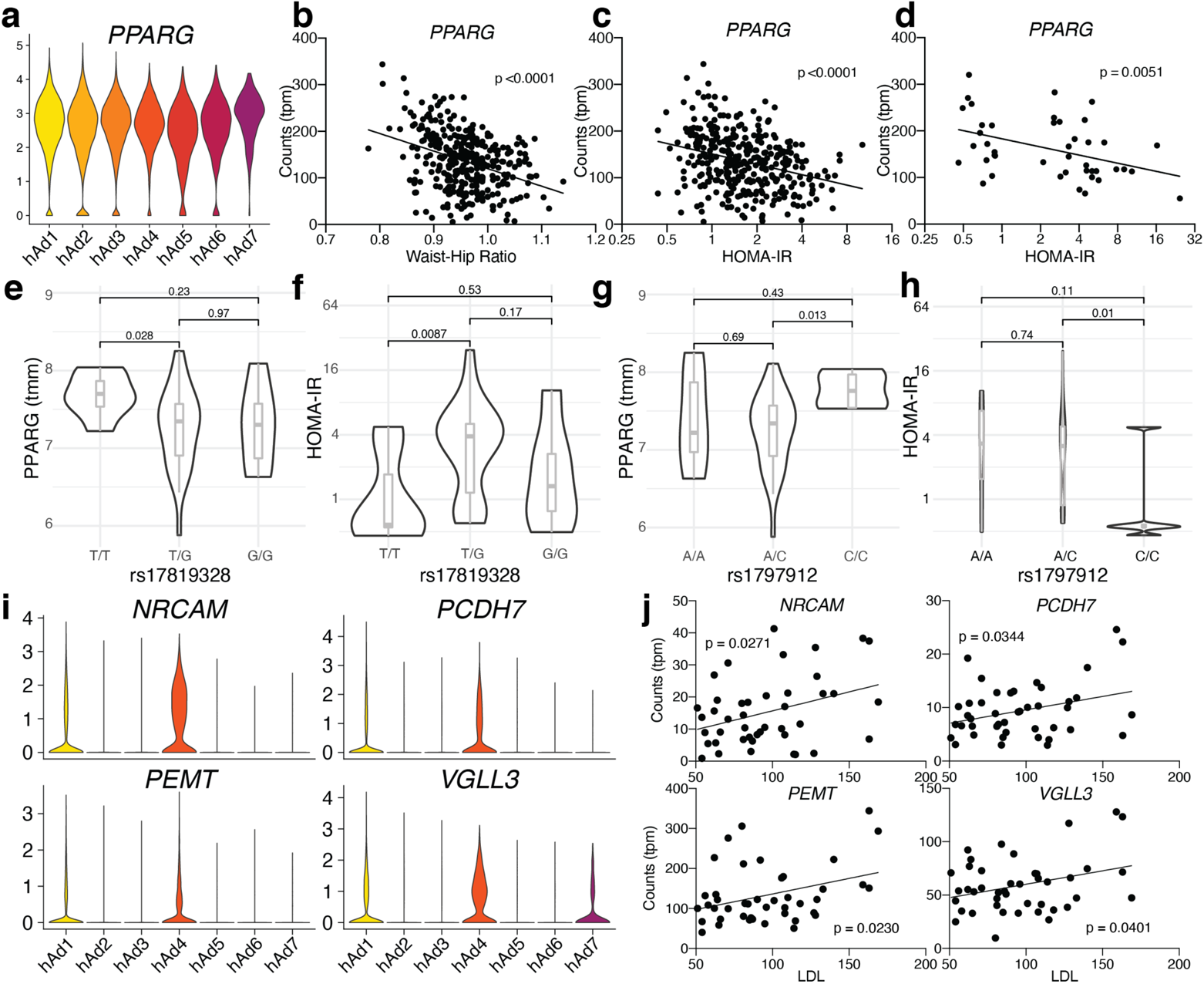
Association with GWAS data provides further insight into the contribution of human white adipocytes to human traits. **a-c,** Expression of *PPARG* in human adipocyte subclusters (**a**), and in METSIM SAT bulk RNA-seq plotted against WHR (**b**) or HOMA-IR (**c**). **d,** Expression of *PPARG* in isolated subcutaneous adipocyte bulk RNA-seq plotted against HOMA-IR. **e-h,** SNPs in the *PPARG* gene identified by DEPICT as associated with BMI-adjusted WHR plotted against *PPARG* gene expression (**e, g**) and HOMA-IR (**f, h**) in isolated subcutaneous adipocyte bulk RNA-seq data and cohort. **i-j,** Expression of genes in human adipocyte subtypes from sNuc-seq data (**i**) and from isolated subcutaneous adipocyte bulk RNA-seq plotted against LDL levels (**j**).

**Extended Data Fig. 18.**
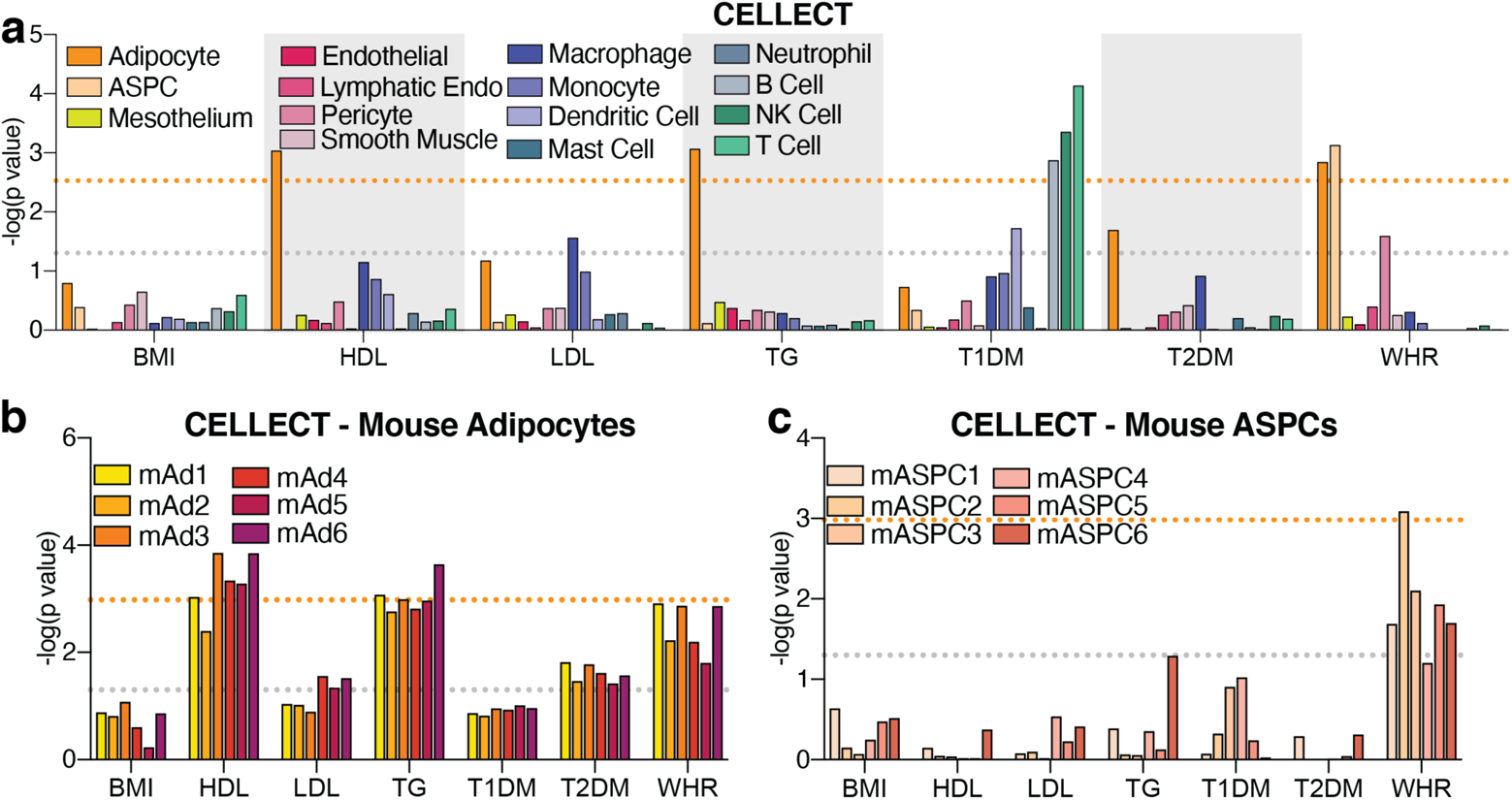
CELLECT identifies mouse cell types associated with human GWAS studies. **a,** *p* values of the association between mouse cell types and GWAS studies. **b-c,** *p* values of the association between mouse adipocyte (**b**) or ASPC (**c**) subclusters with GWAS studies. For all graphs, the grey line represents *p* = 0.05 and the orange line represents significant *p* value after Bonferroni adjustment (*p* = 0.003 for all cell, *p* = 0.001 for subclusters), calculated based on number of cell types queried.

**Extended Data Table 1.**
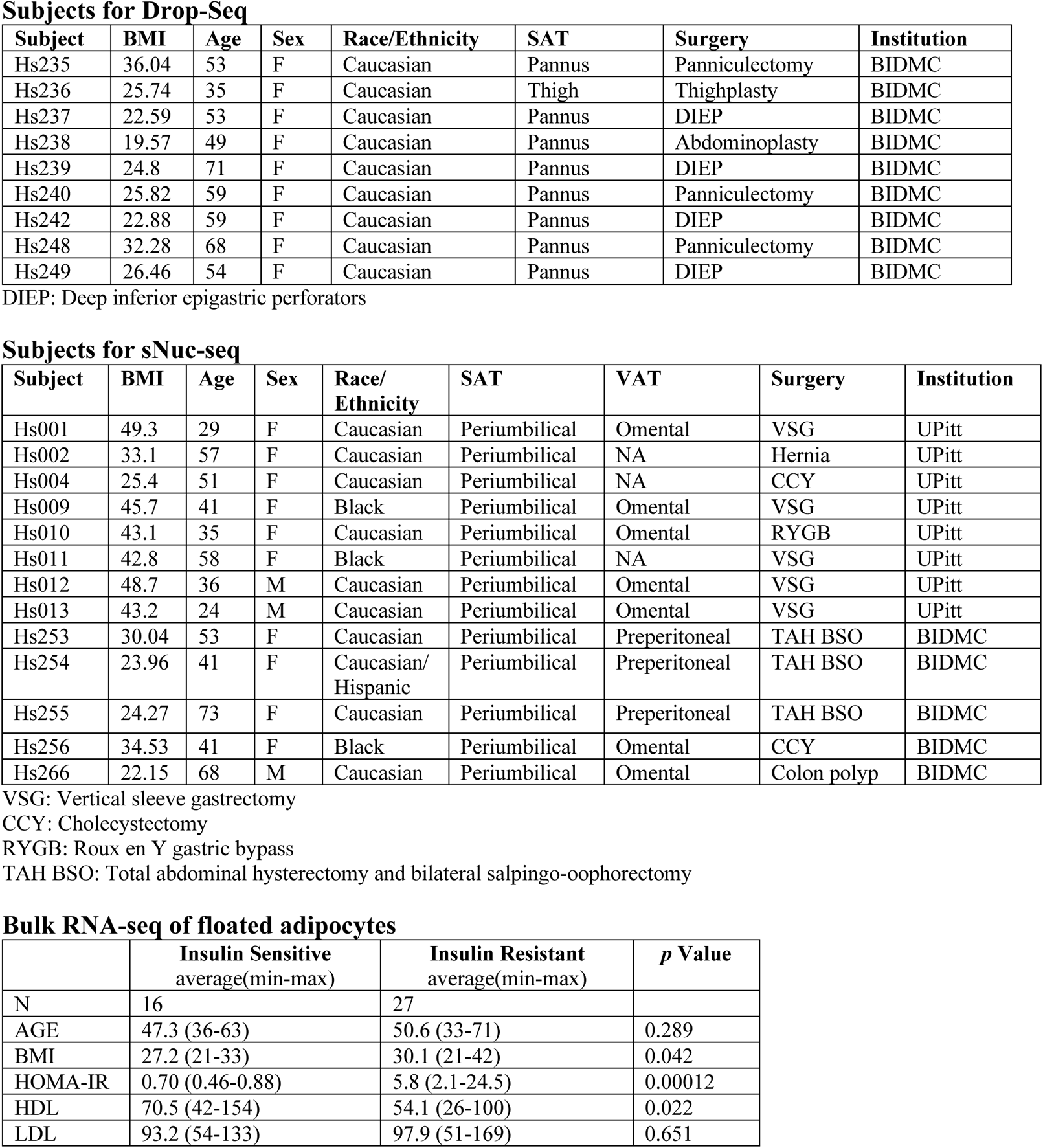
Subject information for Drop-Seq, sNuc-seq, and bulk RNA-seq of isolated subcutaneous human adipocytes

**Subjects for Drop-Seq**
| Subject | BMI | Age | Sex | Race/Ethnicity | SAT | Surgery | Institution |
| --- | --- | --- | --- | --- | --- | --- | --- |
| Hs235 | 36.04 | 53 | F | Caucasian | Pannus | Panniculectomy | BIDMC |
| Hs236 | 25.74 | 35 | F | Caucasian | Thigh | Thighplasty | BIDMC |
| Hs237 | 22.59 | 53 | F | Caucasian | Pannus | DIEP | BIDMC |
| Hs238 | 19.57 | 49 | F | Caucasian | Pannus | Abdominoplasty | BIDMC |
| Hs239 | 24.8 | 71 | F | Caucasian | Pannus | DIEP | BIDMC |
| Hs240 | 25.82 | 59 | F | Caucasian | Pannus | Panniculectomy | BIDMC |
| Hs242 | 22.88 | 59 | F | Caucasian | Pannus | DIEP | BIDMC |
| Hs248 | 32.28 | 68 | F | Caucasian | Pannus | Panniculectomy | BIDMC |
| Hs249 | 26.46 | 54 | F | Caucasian | Pannus | DIEP | BIDMC |

DIEP: Deep inferior epigastric perforators

**Subjects for sNuc-seq**
| Subject | BMI | Age | Sex | Race/<br>Ethnicity | SAT | VAT | Surgery | Institution |
| --- | --- | --- | --- | --- | --- | --- | --- | --- |
| Hs001 | 49.3 | 29 | F | Caucasian | Periumbilical | Omental | VSG | UPitt |
| Hs002 | 33.1 | 57 | F | Caucasian | Periumbilical | NA | Hernia | UPitt |
| Hs004 | 25.4 | 51 | F | Caucasian | Periumbilical | NA | CCY | UPitt |
| Hs009 | 45.7 | 41 | F | Black | Periumbilical | Omental | VSG | UPitt |
| Hs010 | 43.1 | 35 | F | Caucasian | Periumbilical | Omental | RYGB | UPitt |
| Hs011 | 42.8 | 58 | F | Black | Periumbilical | NA | VSG | UPitt |
| Hs012 | 48.7 | 36 | M | Caucasian | Periumbilical | Omental | VSG | UPitt |
| Hs013 | 43.2 | 24 | M | Caucasian | Periumbilical | Omental | VSG | UPitt |
| Hs253 | 30.04 | 53 | F | Caucasian | Periumbilical | Preperitoneal | TAH BSO | BIDMC |
| Hs254 | 23.96 | 41 | F | Caucasian/<br>Hispanic | Periumbilical | Preperitoneal | TAH BSO | BIDMC |
| Hs255 | 24.27 | 73 | F | Caucasian | Periumbilical | Preperitoneal | TAH BSO | BIDMC |
| Hs256 | 34.53 | 41 | F | Black | Periumbilical | Omental | CCY | BIDMC |
| Hs266 | 22.15 | 68 | M | Caucasian | Periumbilical | Omental | Colon polyp | BIDMC |

VSG: Vertical sleeve gastrectomy
CCY: Cholecystectomy
RYGB: Roux en Y gastric bypass

**Bulk RNA-seq of floated adipocytes**
|  | Insulin Sensitive<br>average(min-max) | Insulin Resistant<br>average(min-max) | *p* Value |
| --- | --- | --- | --- |
| N | 16 | 27 |  |
| AGE | 47.3 (36-63) | 50.6 (33-71) | 0.289 |
| BMI | 27.2 (21-33) | 30.1 (21-42) | 0.042 |
| HOMA-IR | 0.70 (0.46-0.88) | 5.8 (2.1-24.5) | 0.00012 |
| HDL | 70.5 (42-154) | 54.1 (26-100) | 0.022 |
| LDL | 93.2 (54-133) | 97.9 (51-169) | 0.651 |

**Extended Data Table 2.** Numbers of cells in human and mouse single cell experiments broken down by cluster, depot, BMI/diet, and technology

**Human Cell Numbers**
| BMI | VAT |  |  | SAT |  |  |  |  | VAT<br>total | SAT<br>total | Total |
| --- | --- | --- | --- | --- | --- | --- | --- | --- | --- | --- | --- |
|  | sNuc |  |  | sNuc |  |  | Drop |  |  |  |  |
|  | < 30 | 30-40 | > 40 | < 30 | 30-40 | > 40 | < 30 | > 30 |  |  |  |
| Adipocyte | 5211 | 1011 | 5253 | 7611 | 2847 | 3938 | 0 | 0 | 11475 | 14396 | 25871 |
| ASPCs | 5938 | 1404 | 7304 | 6848 | 2703 | 7329 | 15195 | 5761 | 14646 | 37836 | 52482 |
| Mesothelium | 7773 | 1927 | 20782 | 0 | 0 | 0 | 0 | 0 | 30482 | 0 | 30482 |
| Endothelial | 2351 | 1030 | 2345 | 4231 | 2783 | 2059 | 577 | 107 | 5726 | 9757 | 15483 |
| Lymphatic Endo | 677 | 240 | 1138 | 195 | 130 | 305 | 168 | 48 | 2055 | 846 | 2901 |
| Pericyte | 381 | 109 | 254 | 353 | 132 | 172 | 60 | 3 | 744 | 720 | 1464 |
| Smooth Muscle | 448 | 360 | 423 | 709 | 621 | 237 | 83 | 5 | 1231 | 1655 | 2886 |
| Macrophage | 1908 | 630 | 6328 | 3121 | 1795 | 2871 | 1256 | 403 | 8866 | 9446 | 18312 |
| Monocyte | 98 | 41 | 173 | 187 | 155 | 549 | 359 | 387 | 312 | 1637 | 1949 |
| Dendritic Cell | 125 | 30 | 340 | 169 | 119 | 188 | 756 | 714 | 495 | 1946 | 2441 |
| Mast Cell | 111 | 27 | 139 | 210 | 294 | 298 | 66 | 23 | 277 | 891 | 1168 |
| Neutrophil | 7 | 9 | 4 | 98 | 12 | 14 | 0 | 2 | 20 | 126 | 146 |
| B Cell | 28 | 12 | 39 | 57 | 49 | 188 | 30 | 26 | 79 | 350 | 429 |
| NK Cell | 229 | 92 | 242 | 375 | 279 | 669 | 297 | 446 | 563 | 2066 | 2629 |
| T Cell | 762 | 382 | 1661 | 667 | 510 | 1522 | 977 | 713 | 2805 | 4389 | 7194 |
| Endometrium | 45 | 150 | 114 | 0 | 0 | 0 | 2 | 1 | 309 | 3 | 312 |
| Total | 26092 | 7454 | 46539 | 24831 | 12429 | 20339 | 19826 | 8639 | 80085 | 86064 | 166149 |

**Mouse Cell Numbers**
|  | PG |  | Ing |  | PG Total | Ing Total | Total |
| --- | --- | --- | --- | --- | --- | --- | --- |
|  | Chow | HFD | Chow | HFD |  |  |  |
| Adipocyte | 12874 | 5139 | 8645 | 13276 | 18013 | 21921 | 39934 |
| ASPCs | 9928 | 10194 | 16308 | 14797 | 20122 | 31105 | 51227 |
| Mesothelium | 10074 | 4873 | 0 | 0 | 14947 | 0 | 14947 |
| Endothelial | 1521 | 673 | 1141 | 2261 | 2194 | 3402 | 5596 |
| Lymphatic Endo | 678 | 101 | 224 | 173 | 779 | 397 | 1176 |
| Pericyte | 62 | 170 | 56 | 309 | 232 | 365 | 597 |
| Smooth Muscle | 56 | 52 | 30 | 125 | 108 | 155 | 263 |
| Macrophage | 3788 | 35673 | 9370 | 9017 | 39461 | 18387 | 57848 |
| Monocyte | 975 | 2801 | 1286 | 2545 | 3776 | 3831 | 7607 |
| Dendritic Cell | 268 | 688 | 237 | 379 | 956 | 616 | 1572 |
| Mast Cell | 4 | 267 | 13 | 27 | 271 | 40 | 311 |
| Neutrophil | 23 | 9 | 8 | 7 | 32 | 15 | 47 |
| B Cell | 301 | 594 | 28 | 279 | 895 | 307 | 1202 |
| NK Cell | 110 | 215 | 67 | 282 | 325 | 349 | 674 |
| T Cell | 266 | 472 | 69 | 479 | 738 | 548 | 1286 |
| Male Epithelial | 3463 | 36 | 19 | 329 | 3499 | 348 | 3847 |
| Female Epithelial | 76 | 45 | 6331 | 3135 | 121 | 9466 | 9587 |
| Total | 44467 | 62002 | 43832 | 47420 | 106469 | 91252 | 197721 |

**Extended Data Table 3.** GWAS studies used for CELLECT analysis

| **Trait** | **Study/collection** |
| --- | --- |
| **BMI** | Pulit, S. L. et al. Meta-analysis of genome-wide association studies for body fat distribution in 694 649 individuals of European ancestry. |
| **HDL** | https://alkesgroup.broadinstitute.org/sumstats_formatted/ |
| **LDL** | https://alkesgroup.broadinstitute.org/sumstats_formatted/ |
| **T1D** | https://alkesgroup.broadinstitute.org/sumstats_formatted/ |
| **T2D (BMI adjusted)** | Mahajan, A. et al. Fine-mapping type 2 diabetes loci to single-variant resolution using high-density imputation and islet-specific epigenome maps. |
| **Triglycerides** | https://alkesgroup.broadinstitute.org/sumstats_formatted/ |
| **WHR (BMI adjusted)** | Loh, P.-R., Kichaev, G., Gazal, S., Schoech, A. P. & Price, A. L. Mixed-model association for biobank-scale datasets |

