## Supplemental guide and note for "A single cell atlas of human and mouse white adipose tissue"

SUPPLEMENTARY INFORMATION

**SUPPLEMENTARY INFORMATION GUIDE**

**Supplementary Table 1. Markers for human clusters and subclusters**

**Supplementary Table 2. Markers for mouse clusters and subclusters**

**Supplementary Table 3. GO and KEGG analysis of markers of human and mouse adipocyte subtypes.**

**Supplementary Table 4. Significant interactions identified by CellphoneDB in human and mouse adipose tissue**

**Supplementary Table 5. Average expression of genes in clusters split by high or low BMI (human) or diet (mouse) for genes in interactions identified by CellphoneDB.**

**Supplementary Table 6. Interactions between adipocytes and endothelial cells, ASPCs, and macrophages in human and mouse adipose tissue. TRUE refers to an interaction that is statistically significant under the given condition.**

**Supplementary Table 7. CELLECT output for human and mouse clusters and subclusters.**

**SUPPLEMENTARY NOTES**

**Supplementary Note 1.** The nomenclature of the Lin- fraction of the SVF is complicated and inconsistent. Traditionally they are defined by exclusion, i.e., they are delineated as SVF cells that are not endothelial (*CD31^+^*), immune (*CD45^+^*), or red blood cells (*TER119^+^*). We would add that they are also not smooth muscle cells (*MYOCD^+^*) or pericytes (*RGS5^+^*), both of which are preferentially associated with the vasculature, or mesothelial cells (*MSLN^+^*), which form a distinct layer on the exterior of visceral depots. Lin- cells are fibroblastic in appearance and some serve as stem cells or progenitors for adipocytes. Various authors have referred to these cells as “fibro-adipogenic precursor cells” (FAPs), “ASCs” (which can stand for adipose stromal cells or adipose stem cells), “APCs” (which can stand for “adipose precursor cells” or “adipose progenitor cells”), or pre-adipocytes, among other terms^84,85^. All of these terms are problematic, in that some of these cells may serve a supporting role only and may not be in the direct adipocyte lineage^3^. One could sidestep the issue by focusing on the specific marker genes that delineate each population, but there has been little consensus in the field here as well. There has been a recent attempt to unify the nomenclature, with the term ASPC (“adipose stem and progenitor cells”) chosen as the parent term, with a hierarchy of component cell types . In this paper we have adopted ASPC as our overall term for these Lin- cells.
